# The tryptophan-binding pockets of Arabidopsis AGO1 facilitate amplified RNA interference via SGS3

**DOI:** 10.1101/2025.10.18.683247

**Authors:** Diego López-Márquez, Laura Arribas-Hernández, Christian Poulsen, Emilie Duus Oksbjerg, Nathalie Bouteiller, Mads Meier, Johannes Blanke, Angel del Espino, Maria Louisa Vigh, Simon Bressendorff, Alberto Carbonell, Rune Daucke, Erwin M. Schoof, Hervé Vaucheret, Peter Brodersen

## Abstract

ARGONAUTE (AGO) proteins associate with small RNAs to form RNA-induced silencing complexes (RISCs). Arabidopsis AGO1 effects post-transcriptional silencing by microRNAs (miRNAs) and small interfering RNAs (siRNAs) and is necessary for siRNA amplification through conversion of RISC target RNAs into double-stranded RNA by the RNA-dependent RNA Polymerase RDR6 and its mandatory cofactors SGS3 and SDE5. Many AGO proteins harbor hydrophobic pockets that interact with tryptophan residues, often surrounded by glycine (GW/WG), in intrinsically disordered regions (IDRs) of RISC cofactors. Here, we show that GW/WG dipeptides in the IDR of SGS3 and the hydrophobic pockets in AGO1 are required for fully functional RDR6-dependent siRNA amplification. We also show that this mechanism requires AGO1-specific structural elements, including positively charged residues surrounding the binding pockets, and a conserved, negatively charged patch in the IDR of SGS3. Thus, the same, conserved protein-protein interaction site is used for different purposes in distinct eukaryotic AGO proteins: the GW/WG-mediated TNRC6-Ago2 interaction is crucial for miRNA-guided silencing in metazoans whereas the GW/WG-mediated SGS3-AGO1 interaction facilitates siRNA amplification via RDR6 in plants.

## INTRODUCTION

Small non-coding RNAs including microRNAs (miRNAs) and short interfering RNAs (siRNAs) are key regulators of eukaryotic gene expression. They use base pairing to guide repressive RNA-induced silencing complexes (RISCs) to nucleic acids with sufficient complementarity to the small RNA to cause stable binding of RISC (Zamore & Haley, 2005). The minimal RISC consists of an ARGONAUTE (AGO) protein bound to the small guide RNA (Rivas *et al*, 2005) with 5’- and 3’-ends buried in dedicated binding pockets (Ma *et al*, 2004; Ma *et al*, 2005; Parker *et al*, 2005; Schirle & MacRae, 2012). The 5’-part of the small RNA adopts a helical conformation with Watson-Crick edges of the nucleobases pointing outwards, thus primed for base pairing with potential target RNAs (Schirle & MacRae, 2012; Wang *et al*, 2009).

In addition to small RNA binding, two biochemical properties are recurrent in many eukaryotic AGO proteins. (i) They possess endonuclease activity via a two-metal ion catalytic site in the so-called Piwi-domain that is structurally related to Ribonuclease H (Liu *et al*, 2004; Meister *et al*, 2004; Nakanishi *et al*, 2012; Song *et al*, 2004). For instance, the RNAseH-like activity of Arabidopsis AGO1 (AtAGO1) is responsible for site-specific cleavage of target RNA (“slicing”) that is required for plant miRNA function (Arribas-Hernandez *et al*, 2016a; Baumberger & Baulcombe, 2005; Qi *et al*, 2005). (ii) They harbor hydrophobic pockets that serve as binding sites for indole rings of tryptophan (Trp) side chains in intrinsically disordered regions (IDRs) of RISC co-factors. The Trp residues implicated in AGO binding via interaction with the hydrophobic pockets are often, but not always, surrounded by glycine (El-Shami *et al*, 2007; Pfaff *et al*, 2013; Till *et al*, 2007), and the AGO-binding sites are, consequently, referred to as “GW-binding sites”. Initial structural studies of human Ago2 (HsAgo2) defined two hydrophobic pockets in the Piwi domain as tryptophan binding sites (Schirle & MacRae, 2012). These pockets, referred to here as Pockets 1 and 2, were suggested to be sites of interaction with the GW-repeat protein TNRC6/GW182 (Schirle & MacRae, 2012) that mediates many effects of metazoan miRNAs on target mRNA translatability and degradation (Braun *et al*, 2013). Indeed, subsequent nuclear magnetic resonance (NMR) and crystallographic studies of the TNRC6 interaction with HsAgo2 and HsAgo1 confirmed that Pockets 1 and 2 act as binding sites for indole rings in specific Trp residues in TNRC6 (Elkayam *et al*, 2017; Pfaff *et al*., 2013; Sheu-Gruttadauria & MacRae, 2018), and also uncovered a third Trp-binding site of importance for TNRC6 interaction (Sheu-Gruttadauria & MacRae, 2018), referred to here as Pocket 3.

In plants, the AGO proteins involved in RNA-directed DNA methylation (RdDM), AGO4, AGO6 and AGO9, interact with GW-repeats of the large subunit of RNA Polymerase V and its cognate SPT5-like elongation factor (Bies-Etheve *et al*, 2009; El-Shami *et al*., 2007; Lahmy *et al*, 2016). This interaction is presumed to involve GW-binding pockets in these three AGO proteins, although formal proof by mutational analysis of conserved residues in the pockets has not been conducted. AGO1 is the key factor involved in post-transcriptional gene regulation via miRNAs and many siRNAs in plants (Vaucheret, 2008). The slicer activity of AGO1 is required for most of its functions in miRNA-controlled development (Arribas-Hernandez *et al*., 2016a; Carbonell *et al*, 2012), but AGO1 also has functions in miRNA-mediated translational repression (Brodersen *et al*, 2008) and in the process known as siRNA amplification (Arribas-Hernandez *et al*, 2016b; Fagard *et al*, 2000; Morel *et al*, 2002).

In siRNA amplification, AGO1 loaded with a primary small RNA triggers recruitment of the RNA-dependent RNA polymerase RDR6 to the target RNA to yield double-stranded RNA (dsRNA) for processing into the first wave of amplified siRNAs, the so-called secondary siRNAs. If the target RNA is of foreign nature, for instance virus- or transgene-derived, secondary siRNAs may themselves trigger new waves of siRNAs, thereby establishing a potent siRNA amplification positive feedback loop that is crucial in antiviral defense (Dalmay *et al*, 2000; Garcia-Ruiz *et al*, 2010; Mourrain *et al*, 2000; Wang *et al*, 2011). Plants have also evolved a controlled version of siRNA amplification, the *trans-*acting or phased siRNA (tasiRNA or phasiRNA) pathway, for endogenous gene control. In this pathway, a miRNA-programmed RISC (miRISC) targets a precursor transcript to yield a single wave of secondary siRNAs known as tasiRNAs or phasiRNAs (Liu *et al*, 2020). miRNA-induced secondary siRNAs are of tremendous biological importance, as they are key to morphogenesis, defence against pathogenic fungi and oomycetes, and, in monocots, reproductive development and hence food production (Liu *et al*., 2020). In addition, the fact that tasiRNA biogenesis initiates at a well-defined site – the cleavage site guided by the primary miRNA trigger – makes this pathway well-suited to study mechanistic aspects of the initiation step of siRNA amplification.

At least two distinct settings may instigate secondary siRNA formation. First, if RISC is loaded with a 22-nt miRNA trigger, a single target site is sufficient to trigger amplification (Chen *et al*, 2010; Cuperus *et al*, 2010; Fei *et al*, 2018) and this process may be further promoted if the 22-nt trigger miRNA derives from a bulged 22-nt/21-nt miRNA/miRNA* duplex (Manavella *et al*, 2012). Second, if the target transcript has at least two small RNA target sites, 21-nt miRNAs can also trigger secondary siRNAs through cooperative RISC action whose molecular basis remains ill-defined (Axtell *et al*, 2006; Howell *et al*, 2007). Multiple lines of evidence indicate that RISC itself, not the target RNA cleavage fragments released by slicing as proposed originally (Yoshikawa *et al*, 2005), is the trigger of secondary siRNA formation. For example, catalytically inactive AGO1 loaded with the 22-nt miR173 efficiently induces tasiRNA production (Arribas-Hernandez *et al*., 2016b), and a cleavage-competent RISC consisting of AGO1 and a 21-nt miR173 version fails to induce secondary siRNAs, as do cleavage-competent AGO1-based 21-nt miRISCs in general (Chen *et al*., 2010; Cuperus *et al*., 2010). In addition, biochemical analyses in cell-free lysates indicate that an amplifier complex consisting of the proteins SDE5 and SGS3 in addition to AGO1 loaded with a 22-nt miRNA acts as a platform onto which RDR6 can associate and initiate dsRNA synthesis using the RNA targeted by AGO1 as template (Sakurai *et al*, 2021; Yoshikawa *et al*, 2021). Following completion of dsRNA synthesis, DICER-LIKE4 (DCL4) produces 21-nt siRNA duplexes (Allen *et al*, 2005; Dunoyer *et al*, 2005; Gasciolli *et al*, 2005; Xie *et al*, 2005; Yoshikawa *et al*., 2005), a reaction that also involves the generation of 2-nt 3’-overhangs on the dsRNA by the terminal uridyl transferase (TNTase) HESO1 (Vigh *et al*, 2024).

Although the evidence for a RISC trigger model (Branscheid *et al*, 2015) where AGO1 acts as a scaffold to assemble the AGO1-SGS3-SDE5-RDR6 amplifier complex on target RNA is now considerable, important predictions of this model have not been verified experimentally. Specific interactions between AGO1, SGS3, SDE5 and RDR6 have not been described, and, crucially, mutant alleles of the different components with specific mutual interaction defects have not been isolated and characterized for secondary siRNA production. Several mutants of AGO1 with defects in siRNA amplification have been identified in forward screens (Fagard *et al*., 2000; Morel *et al*., 2002; Poulsen *et al*, 2013), but they all have more general defects in miRNA-guided silencing, presumably due to impaired small RNA binding (Poulsen *et al*., 2013) or siRNA duplex unwinding (Derrien *et al*, 2018). Thus, molecular properties of AGO1 required specifically for small RNA-induced siRNA amplification, including determinants of assembly of the SGS3-AGO1-SDE5 amplifier platform, remain unknown.

At least one GW-binding site in AGO1 is clearly conserved (Poulsen *et al*., 2013), and it is, therefore, a question of outstanding interest what purpose this binding site fulfills. In addition, because plants encode AGO proteins with widely different functions, such as AGO4 in RdDM and AGO1 in miRNA-and siRNA-mediated post-transcriptional silencing (Fang & Qi, 2016), it is also a question of general importance which features, in addition to accommodation of indole side chains, contribute to specificity in AGO-GW interactions.

Here, we construct structure-guided point mutants targeting the conserved GW-binding pockets in AGO1 as well as mutants in two structural elements specific to AGO1 in the vicinity of the GW-binding pockets: a positively charged surface immediately surrounding the pockets, and an insertion of 13 amino acid residues not found in well-studied animal AGOs. In the cryoEM structure of the closest AGO1 paralogue, AGO10, this part adopts a small β-sheet projecting away from the main body of the protein into the solvent and is, therefore, referred to as the β-finger (Xiao *et al*, 2023). We show that inactivation of pockets 1 and 3, as well as mutation of surrounding positively charged residues or deletion of the β-finger leads to clear defects in siRNA amplification, but not in miRNA-guided slicing. Remarkably, mutation of GW/WG dipeptides or a highly conserved negatively charged region in the IDR of SGS3 causes similar defects in siRNA amplification. Hence, we propose that a GW/WG-mediated AGO1-SGS3 interaction facilitates siRNA amplification via RDR6 in plants.

## RESULTS

### Design of AGO1 mutants with predicted defects in GW binding pockets

To design mutants in GW-binding sites in AtAGO1, we first inspected crystal structures of HsAgo2 and alignments of AtAGO1 and HsAgo2 sequences to see if residues integral to function of GW-binding pockets in HsAgo2 were conserved in AtAGO1. This analysis showed that pockets 1 and 3 were conserved, while pocket 2 was self-filled by a Trp residue (W785) not present in HsAgo2 (**Fig. 1 A-C**), consistent with a recent analysis involving both plant and non-plant eukaryotic AGO protein sequences (Wallmann & Van de Pette, 2024). The TNRC6 interaction with HsAgo2 has previously been shown to be abolished in the double mutant K660A/L694Y (corresponding to K839A/L873Y in AtAGO1) (Kuzuoğlu-Öztürk *et al*, 2016). We therefore constructed K839A and L873Y mutants to affect pocket 1 function in two different ways. We also constructed the R867S mutant to abrogate pocket 3 function, as done previously in HsAgo2 (Sheu-Gruttadauria & MacRae, 2018). Finally, we included the F832A mutant whose analogue in fly Ago1 (F777A) affects binding of GW182 (Eulalio *et al*, 2009; Eulalio *et al*, 2008), potentially by interference with the function of pocket 2. These mutants constitute a “GW tester set” to query the function of GW pocket function in AtAGO1 (henceforth AGO1) *in vivo*.

**Figure 1.**
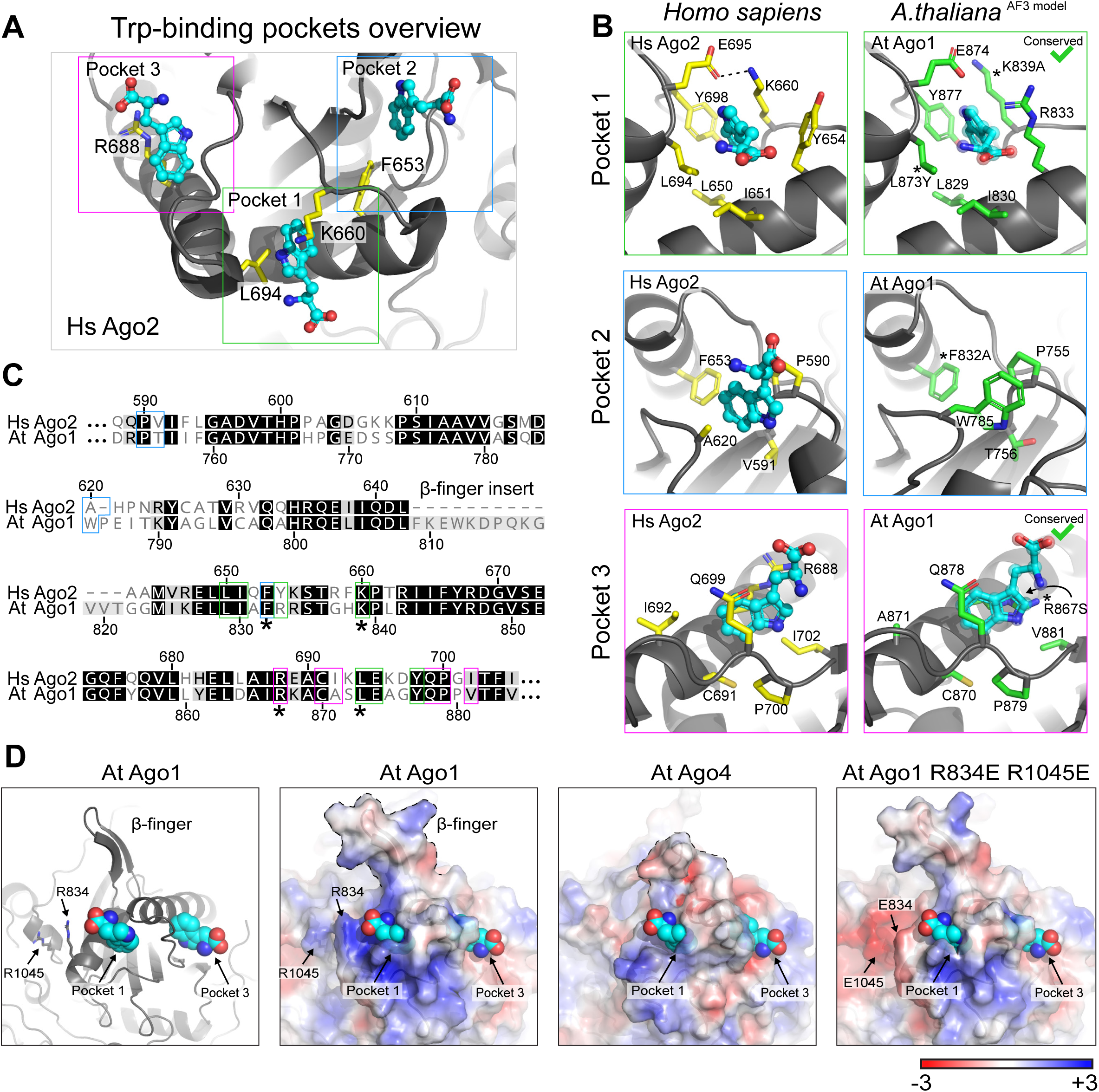
Structural overview of pocket regions in HsAgo2 and AtAGO1. **(A)** Structural model of HsAgo2 showing pocket 1 (green square), pocket 2 (blue square), and pocket 3 (magenta square). Key residues are indicated: L694 and K660 (pocket 1), F653 (pocket 2), and R688 (pocket 3). Tryptophan residues accommodated in the pockets are shown in cyan. **(B)** Structural overview of pockets 1, 2, and 3 in HsAgo2 (PDB: 6CBD) (Sheu-Gruttadauria & MacRae, 2018) and AtAGO1 (Alphafold3 model). Key residues are indicated in yellow (HsAgo2) and green (AtAGO1). Tryptophan residues accommodated in the pockets are shown in cyan. **(C)** Alignment of amino acid sequences corresponding to the pocket regions of HsAgo2 and AtAGO1. Key residues are highlighted in green (pocket 1), blue (pocket 2), and magenta (pocket 3). Black background, identical residues; grey background, conservative changes. Amino acid positions are indicated above and below the sequences. **(D)** Alphafold3 models of the pocket/β-finger region in AtAGO1 and AtAGO4. Charge is indicated in a red (negative) to blue (positive) scale. Residues contributing to the positively charged patch in AGO1 (R834 and R1045) are highlighted. For comparison, the charge distribution of AtAGO4 is shown. A predicted structural model of the AtAGO1 R834E/R1045E mutant is also presented.

### Design of AGO1 mutants with altered features around GW pockets

We next used the Alphafold3-predicted structures of plant AGO proteins, supported by the recently published 3.6Å cryoEM structures of AtAGO10 (Xiao *et al*., 2023), to analyze potential differences between functionally different AGO paralogues. Comparison between AGO1 and AGO proteins in the AGO4/6/9 clade showed a conspicuous difference in the surface surrounding the GW-binding pockets. AGO1 has positively charged amino acids concentrated in this region, while AGO4 has a mosaic of negatively and positively charged residues (**Fig. 1D**). For this reason, we also mutated the AGO1-specific surface-exposed arginine residues R834 and R1045 to glutamate (R834E/R1045E, RR-EE) to change the charge distribution around GW pockets (**Fig. 1D and EV1**). We further noticed that AGO proteins of the AGO1/5/10 and AGO4/6/9 clades harbor an 8-13 amino acid insertion in the GW-pocket binding area compared to animal AGOs and to the AGO2/3/7 clade (**Fig. EV1**). The AGO1 insertion is deeply conserved across orthologues and forms a conspicuous structural element termed the “β-finger” (**Fig. 1C,D and Fig. EV1**) (Xiao *et al*., 2023). We therefore also included a deletion mutant of the β-finger in our analyses.

### AGO1 pocket mutants complement developmental defects of ago1 null mutants

As a rapid test for miRNA function, we scored the ability of the AGO1 mutants to complement the strong developmental phenotype of the *ago1-3* null mutant (Bohmert *et al*, 1998). All mutants were expressed as N-terminally FLAG-tagged versions to facilitate subsequent molecular analyses involving immuno-affinity purification. No interference of the N-terminal FLAG tag with wild type AGO1 function was noted in previous analyses (Arribas-Hernandez *et al*., 2016a; Arribas-Hernandez *et al*., 2016b). We found that all GW pocket-related mutants fully complemented developmental phenotypes of the *ago1-3* mutant (**Fig. 2A,B**), suggesting that the GW pocket mutations do not interfere substantially, if at all, with small RNA binding or repression of developmentally important miRNA targets. Indeed, immuno-affinity purified AGO1 showed similar amounts of miRNAs associated with FLAG-AGO1^WT^ and the GW tester set of FLAG-AGO1 mutants (**Fig. 2C**), corroborating the conclusion that miRNA association is unaffected by pocket mutation.

**Figure 2.**
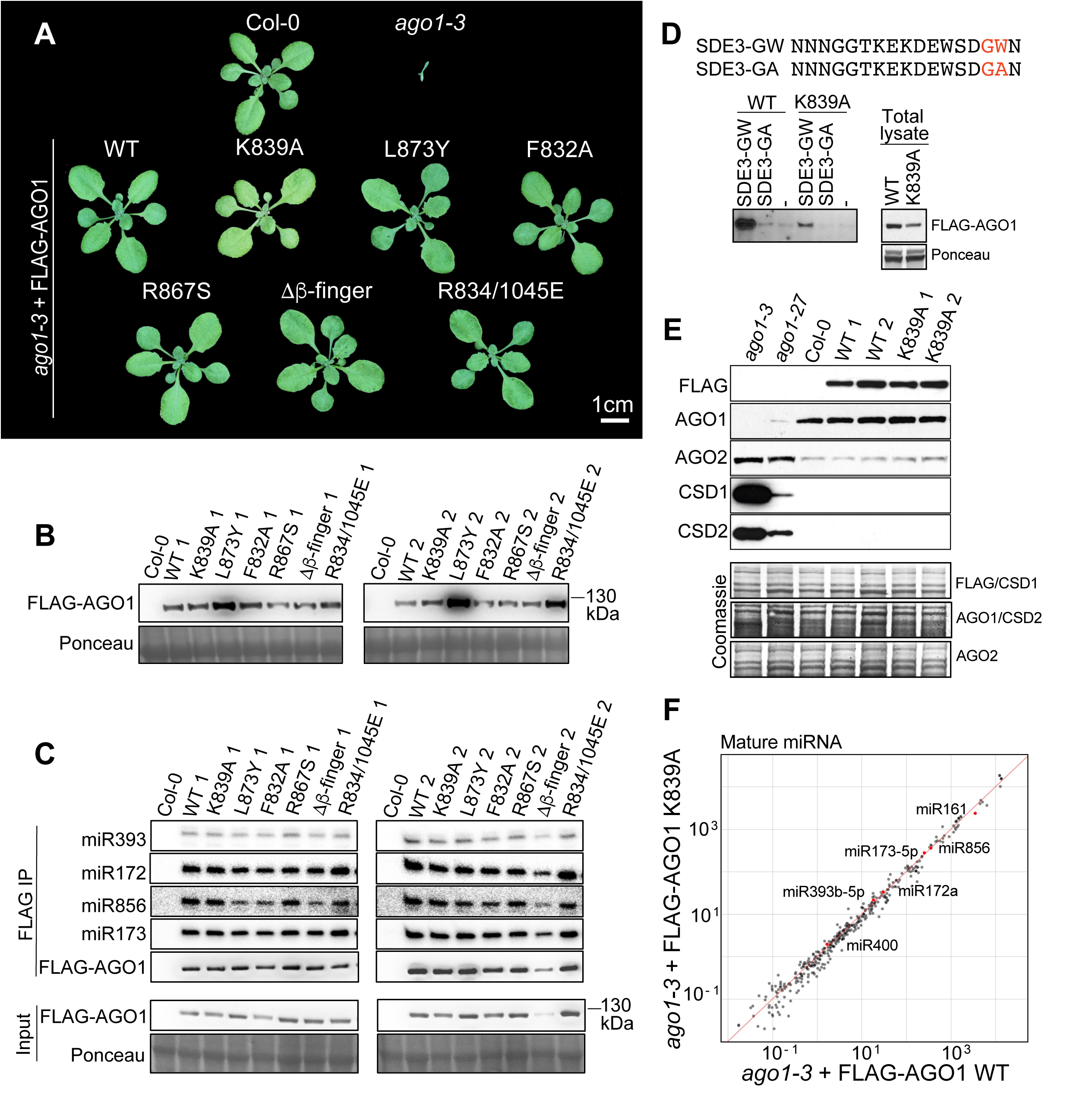
Disruption of tryptophan-binding pockets or adjacent structural elements in AGO1 does not impair miRNA activity. **(A)** Rosettes of 3.5-week-old Arabidopsis plants of the indicated genotypes. **(B)** Immunoblot analysis of total protein extracts from seedling tissue of the genotypes described in (A). Ponceau staining was used as a loading control. **(C)** RNA blot analysis of miRNAs associated with immunoprecipitated FLAG-AGO1 (FLAG IP) from inflorescence tissue of two independent transgenic lines (L1, L2) of each of the FLAG-AGO1 variants shown in (A). The same membrane was sequentially hybridized with different probes. The lower panel shows detection of the FLAG-AGO1 variants in total extracts by protein blot (input). Ponceau staining was used as a loading control. **(D)** In vitro binding assay using total inflorescence lysates prepared from FLAG-AGO1^WT^ or FLAG-AGO1^K839A^-expressing plants and SDE3-derived biotinylated GW (wild type) and GA (mutant) peptides immobilized on streptavidin-conjugated beads. Left panel, bound fractions corresponding to proteins retained by the immobilized peptides. Right panel, AGO1 levels in total protein lysates used as input for the assay. Ponceau staining was used as a loading control for the input samples. **(E)** Protein blot analyses of known AGO1 targets (AGO2, CSD1, and CSD2) in seedlings of wild type Col-0, *ago1* mutants (*ago1-3* and *ago1-27*), and two independent transgenic lines (L1, L2) expressing either FLAG-AGO1^WT^ or FLAG-AGO1^K839A^ in the *ago1-3* background. The anti-AGO1 antibody is used to compare the expression of FLAG-AGO1 transgenes to that of endogenous AGO1 in non-transgenic Col-0 wild type plants. Coomassie staining was used as a loading control. **(F)** Scatter plot showing log₁₀-transformed reads per million (RPM) values of individual miRNAs detected in *ago1-3* plants complemented with either FLAG-AGO1^WT^ or FLAG-AGO1^K839A^. miRNAs of particular relevance to this study are highlighted in red. The red diagonal line represents equal abundance between both genotypes.

### The AGO1^K839A^ mutant exhibits defective GW-peptide interaction

We next used a peptide-based pull-down binding assay to verify the predicted defects in GW-peptide binding of AGO1 pocket inactivation. We chose the K839A pocket 1 mutant and used a GW-containing peptide from the C-terminal IDR of the SILENCING-DEFICIENT3 (SDE3) helicase (El-Shami *et al*., 2007; Garcia *et al*, 2012), because SDE3 has previously been shown to associate with AGO1 dependent on GW dipeptides in its C-terminal IDR (Garcia *et al*., 2012). The results showed that the AGO1^K839A^ mutant protein was retained substantially less efficiently than AGO1^WT^ from total lysates by the SDE3-GW peptide, and that binding was abolished by W-A mutation in the SDE3 peptide (**Fig. 2D**). We note that a partial reduction in GW-peptide interaction is the expected outcome of this test given the presence of the conserved pocket 3 in AGO1. We conclude that pocket inactivation, at least in the case of the AGO1^K839A^ mutant, confers the expected biochemical defect in GW-peptide interaction.

### AGO1 pocket mutants show no apparent defects in miRNA-guided translational repression

We next tested whether defects in miRNA-guided translational repression could be detected. Several studies have shown that repression of COPPER/ZINC SUPEROXIDE DISMUTASE1/2 (CSD1/2) by miR398 involves AGO1-mediated translational repression (Brodersen *et al*., 2008; Li *et al*, 2013). We therefore measured CSD1 and CSD2 protein levels in the K839A (pocket 1) mutant. No differences in CSD1/2 protein accumulation were found compared to wild type, unlike in the hypomorphic *ago1-27* or null *ago1-3* mutants (**Fig. 2E**). Similar results were found for AGO2 whose mRNA is targeted by miR403 (**Fig. 2E**). We conclude that GW pockets of AGO1 are unlikely to be involved in miRNA-guided translational repression. Finally, we performed small RNA-seq of all AGO-bound small RNAs (Grentzinger *et al*, 2020) to avoid taking confounding mRNA degradation fragments into account. This analysis revealed no major differences in miRNA profiles between the K839A pocket 1 mutant and WT plants (**Fig. 2F**).

### GW-binding sites are required for normal siRNA accumulation from many loci

We next tested whether GW-binding sites may be involved in AGO1-dependent secondary siRNA production. Initial analyses by northern blots showed that the tasiRNA siR255 triggered by the AGO1-associated 22-nt miR173 accumulated normally in mutants defective in GW binding sites (**Fig. 3A**). This suggests that GW-binding sites are not universally required for secondary siRNA production. On the other hand, the abundance of *AFB2* secondary siRNAs triggered by the 22-nt miRNA miR393 was reduced in the K839A, RR-EE and Δβ-finger mutants compared to wild type (**Fig. 3A**). We therefore performed small RNA-seq, using the same AGO-enrichment approach as above, to clarify the extent to which the requirement of GW-binding pockets for secondary siRNA production was widespread. All mutants exhibited total miRNA levels, small RNA size-distributions, and 5’ nucleotide compositions comparable to those observed in FLAG-AGO1 WT (**Fig. EV2**). Interestingly, however, independent transgenic lines of several mutants – among them those affecting pockets 1 and 3 – showed clearly differentially expressed siRNA populations (**Fig. EV3A**), including the *AFB2* siRNA population seen by RNA blots (**Fig. 3B**). The small RNA-seq analysis also confirmed that tasiRNA populations induced by miR173-AGO1 were unaffected in the AGO1 pocket mutants (**Fig. 3B**), and that no siRNA populations with significantly different abundance compared to wild type could be identified in the F832A mutant (pocket 2). We conclude from these initial mutational analyses that no functional defects could be identified in the F832A mutant, consistent with the conclusion that pocket 2 does not exist as a functional interaction site in plant AGO1. In contrast, differentially expressed siRNA populations worthy of further analysis were identified in pocket 1 and pocket 3 mutants.

**Figure 3.**
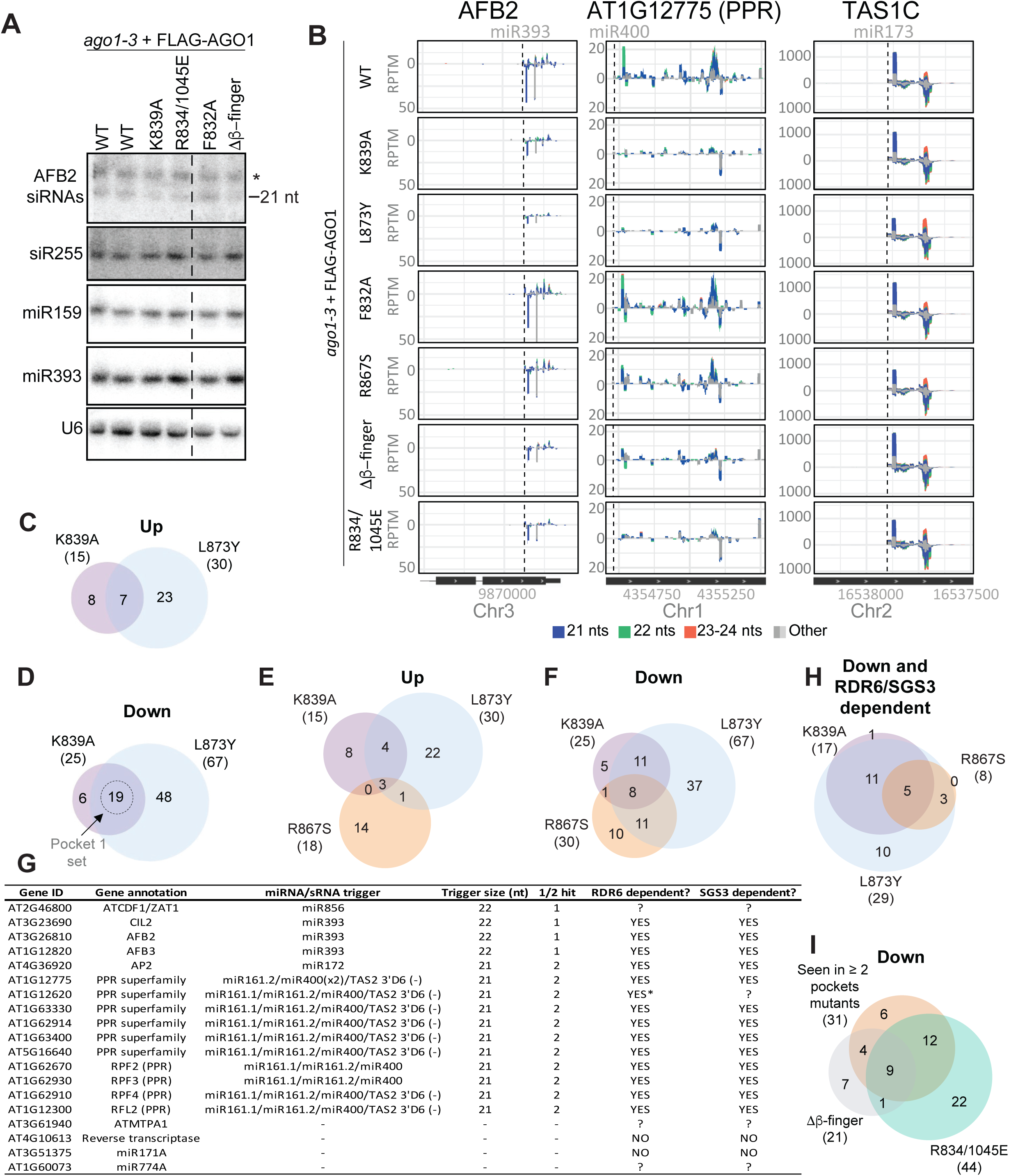
AGO1 pockets and surrounding structural features are required for secondary siRNA production. **(A)** RNA blot analysis of secondary siRNAs (AFB2 and TAS1C), the AFB2 trigger miRNA (miR393), and a control miRNA (miR159) in inflorescences of *ago1-3* plants expressing either the FLAG-AGO1^WT^ or the indicated mutant versions. U6 served as a loading control. The presence of lanes on the original blot cropped for presentation purposes is indicated by a dashed line. The entire blot is provided in Appendix Figure 1. Another part of this blot makes up Figure 2D in the report by (Vigh *et al*., 2024). **(B)** Examples of siRNA accumulation in Reads Per Ten Million (RPTM) for two representative genes (AFB2 and AT1G12775) in inflorescences of *ago1-3* plants expressing either FLAG-AGO1^WT^ or the indicated mutant versions. TAS1C siRNAs are shown as a control. Gene models and genomic coordinates are shown below each plot (note: only partial gene models are displayed). miRNA target sites are highlighted with a dashed line. Small RNAs are color-coded by length: <21 nt (dark grey), 21 nt (blue), 22 nt (green), 23–24 nt (orange), and >24 nt (light grey). **(C)** Venn diagram illustrating the overlap of genes giving rise to siRNAs more abundant in AGO1^K839A^ and AGO1^L873Y^ mutants than in AGO1^WT^ (“Up”). **(D)** Venn diagram illustrating the overlap of genes giving rise to siRNAs less abundant in AGO1^K839A^ and AGO1^L873Y^ mutants than in AGO1^WT^ (“Down”). **(E)-(F)** Venn diagrams illustrating the overlap of upregulated (E) and downregulated (F) siRNA populations between AGO1^K839A^, AGO1^L873Y^ and AGO1^R867S^ mutants. The gene sets were defined by pairwise comparisons to AGO1^WT^. **(G)** Summary of features of the set genes giving rise to less abundant siRNAs in pocket 1 mutants than in AGO1^WT^ (as defined in panel D). An * indicates data from (Branscheid *et al*., 2015); all other data were obtained from libraries generated in this study. **(H)** Venn diagram illustrating the overlap of siRNA populations that are both downregulated in AGO1^K839A^, AGO1^L873Y^ or AGO1^R867S^ mutants compared to AGO1^WT^, and are RDR6/SGS3 dependent. **(I)** Venn diagram showing the overlap in genes giving rise to downregulated siRNA populations between AGO1^pockets^, AGO1^1β-finger^ and AGO1^RR-EE^ mutants. The AGO1^pockets^ set of genes includes the 31 genes with downregulated siRNA populations that overlap between any two of the AGO1^K839A^, AGO1^L873Y^, and AGO1^R867S^ mutants, as illustrated in (F).

### Pockets 1 and 3 are required for many RDR6/SGS3-dependent siRNA populations induced by 22-nt miRNAs or double-hit 21-nt small RNAs

We first divided the differentially expressed small RNA populations into up-regulated and down-regulated groups and analyzed the number of loci affected and the overlap of affected loci between mutants. For the pocket 1 mutants (K839A and L873Y), the loci with reduced small RNA accumulation were more numerous than those with increased accumulation (**Fig. EV3B**). Importantly, the overlap of small RNA-producing loci exhibiting down-regulation between pocket 1 and pocket 3 mutants was clearly larger than that of the upregulated groups (**Fig. 3C-F**). Because of this recurrent defect in small RNA accumulation, we conclude that a primary function of GW-binding pockets in AGO1 is to promote small RNA production and focused on the overlap of 19 loci with reduced small RNA accumulation in both L873A and K839A mutants (**Fig. 3D,G**) for further characterization.

The small RNAs from these 19 loci were predominantly 21 nt in size (**Fig. EV3C**), and did not show any clear 5’-nucleotide bias (**Fig. EV3D**). Since AGO1-associated small RNAs show a strong 5’-U bias (Mi *et al*, 2008; Takeda *et al*, 2008), this observation argues that loss of small RNAs in the pocket mutants is not due to a defect in AGO1 loading, consistent with the largely intact profiles of AGO1-associated miRNAs in the pocket mutants (**Fig. EV2A, E**). Two of the 19 loci were miRNAs (miR171a and miR774a) and were not considered further. Of the remaining 17 loci, 15 were known small RNA targets, featuring either a single target site of a 22-nt miRNA (miR393, miR856), or two or more target sites for 21-nt miRNAs and/or siRNAs (miR172, miR161/miR400/TAS1/TAS2; **Fig. 3G**). In all cases, the affected siRNA populations were adjacent to target sites, consistent with their dependence on a small RNA trigger (**Fig. 3B**). We finally verified their RDR6-and SGS3-dependence, previously described for some of the siRNAs (Howell *et al*., 2007), by small RNA-seq of flowers from *rdr6-12* and *sgs3-1* knockout mutants grown under the same conditions as the AGO1 mutant series (**Fig. 3G-H and Fig. EV4A-F**). This analysis also clearly showed that the defect in production of RDR6-SGS3-dependent siRNAs was recurrent among pocket 1 and pocket 3 mutants, and that the impact of the mutations with respect to number of loci affected decreased in the order L873Y (pocket 1) > K839A (pocket 1) > R867S (pocket 3) (**Fig. 3E-F, H and EV3B**). We conclude that pockets 1 and 3 are required for siRNA amplification triggered by either single target sites of several, but not all, 22-nt miRNAs, or by cooperatively acting 21-nt miRNAs, in both cases via the canonical RDR6-SGS3-dependent pathway.

### A positively charged surface and the β-finger are required for siRNA amplification

Having defined the effects on siRNA amplification of the GW tester set of mutants, we moved on to probe the effects on mutants in the surrounding structural elements: FLAG-AGO1^RR-EE^ and FLAG-AGO1^Δβ-finger^. Several small RNA loci showed markedly reduced abundance in FLAG-AGO1^RR-EE^ and FLAG-AGO1^Δβ-finger^ compared to wild type (**Fig. 3A-B and EV3A-B**). Remarkably, the siRNA-populations affected in FLAG-AGO1^RR-EE^ and FLAG-AGO1^Δβ-finger^ overlapped significantly with those affected in pocket 1 and 3 mutants (**Fig. 3I**), suggesting that a common biochemical defect in these mutants underlies the defective production of RDR6-/SGS3-dependent secondary siRNAs. We conclude that pockets 1 and 3 as well as the β-finger and the positively charged surface surrounding the pockets are required for siRNA amplification, suggesting that binding to co-factors implicated in siRNA amplification involves GW-pocket binding, the β-finger, and a correct charge distribution around the pockets.

### A candidate approach aimed at identification of GW interactors

We next aimed to identify proteins with GW dipeptides that would explain the requirement of Trp-binding pockets in AGO1 for miRNA-induced siRNA production. Differential enrichment analysis between proteins detected by mass spectrometry in FLAG-AGO1^WT^ and FLAG-AGO1^K839A^ immuno-affinity purifications did not identify any candidates (**Fig. EV5A-B**). We therefore took a candidate approach focusing on proteins known to be involved in siRNA amplification. We considered the helicase SDE3 to be an obvious candidate for in-depth genetic analysis for several reasons. It co-immunoprecipitates with AGO1 in a manner dependent on Trp residues in its C-terminal IDR (Garcia *et al*., 2012), and is implicated in some, but notably not all, siRNA amplification phenomena (Dalmay *et al*, 2001; Garcia *et al*., 2012; Himber *et al*, 2003; Jauvion *et al*, 2010; Vazquez *et al*, 2004). For example, SDE3 is necessary for amplified silencing of weak transgenic silencing inducers (Jauvion *et al*., 2010), but is not necessary for tasiRNA generation (Vazquez *et al*., 2004), similar to GW pockets in AGO1. To identify additional candidates, we reasoned that a *bona fide* AGO1 pocket interactor should satisfy three requirements. First, it should harbor conserved Trp residues in IDRs. Second, it should have conserved negatively charged residues that may or may not be in IDRs. Third, mutation of candidate Trp and negatively charged residues should give an siRNA profile similar to that of pocket mutants in AGO1.

We considered as possible candidates the proteins SGS3, SDE5, RDR6 and DCL4 that are required for all miRNA-induced secondary siRNA formation (Allen *et al*., 2005; Jauvion *et al*., 2010; Xie *et al*., 2005; Yoshikawa *et al*., 2005). We also included the family of TNTases (HESO1, URT1, NTP2-9), because HESO1 was recently shown to be involved in secondary siRNA production (Vigh *et al*., 2024), and because TNTases are well known to be implicated in small RNA amplification in *Schizosaccharomyces pombe* and *Caenorhabditis elegans* (Motamedi *et al*, 2004; Shukla *et al*, 2020).

We first used protein disorder prediction software to screen candidate proteins for the presence of Trp residues in IDRs. This filtering step retained SGS3, HESO1, and NTP8 as possible candidates. Of these proteins, only SGS3 and the known GW-interactor SDE3 had clear negatively charged patches in their IDRs (**Fig. EV5C**). We next examined the conservation of Trp residues and the negatively charged patch in IDRs of SGS3 across 196 plant species (Bélanger *et al*, 2023). This analysis revealed that both occurrence of Trp residues and negatively charged residues were highly conserved features (**Fig. 4A-E**). These patterns of sequence conservation qualify SGS3 as an additional candidate for in-depth genetic analyses, further supported by its association with AGO1 in co-immunoprecipitation assays from both intact tissues (**Fig. EV5A-B**) (Yoshikawa *et al*, 2013) and cell-free in vitro translation systems (Sakurai *et al*., 2021; Yoshikawa *et al*., 2021).

**Figure 4.**
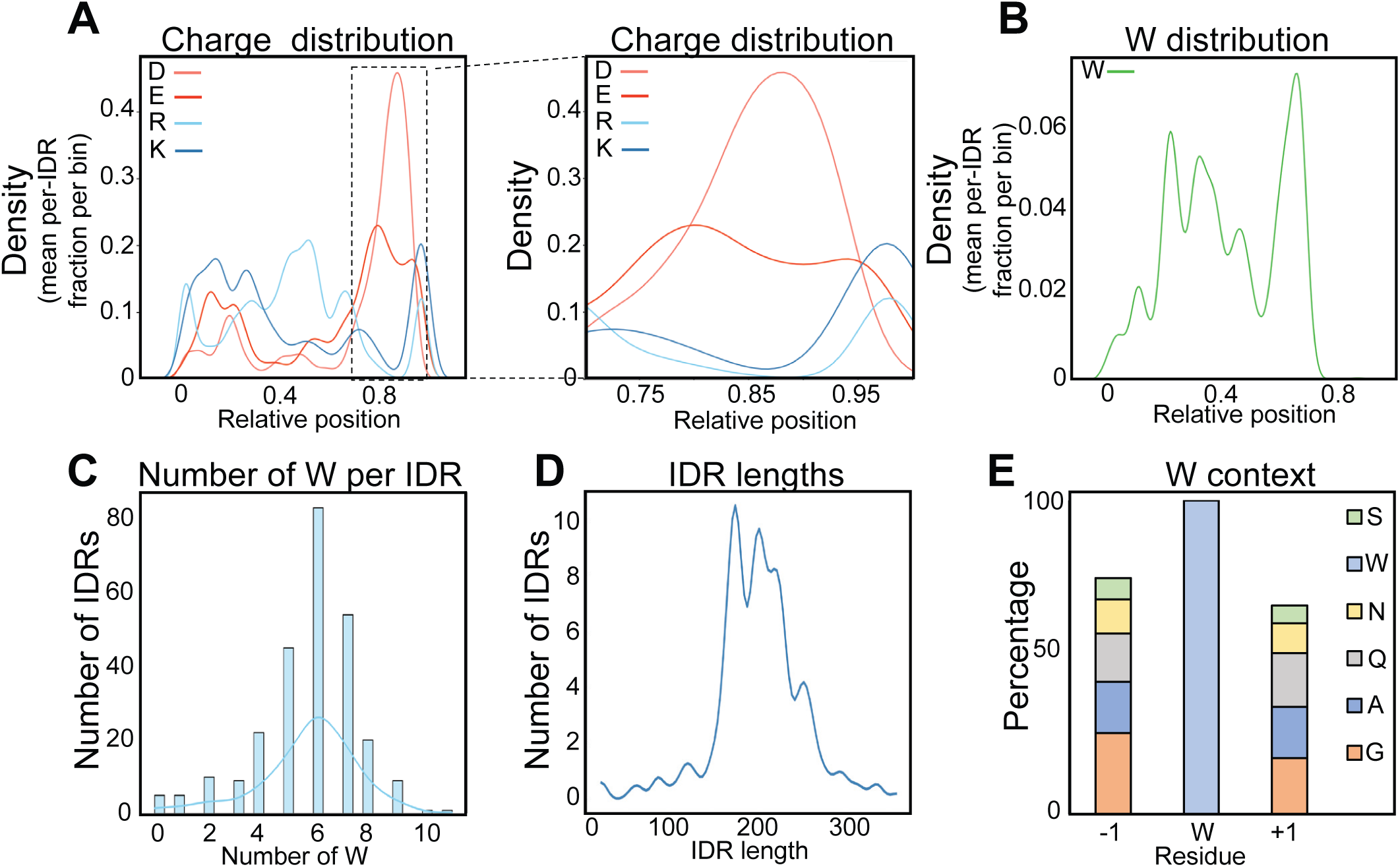
The N-terminal IDR of SGS3 has sequence properties expected for an AGO1 pocket interactor. **(A)** Predicted charge distribution within the SGS3 IDR, based on the frequency of Asp (D), Glu (E), Arg (R), and Lys (K) residues. The analysis was performed using the SGS3 IDR sequence from over 200 plant species (Bélanger *et al*., 2023). A magnified view of a region of special interest is shown on the right. Densities were calculated as the mean fraction per IDR in each bin analyzed. **(B)** Distribution of Trp (W) residues along the SGS3 IDRs analyzed in panel (A). Densities were calculated as in panel (A). **(C)** Histogram depicting the distribution of SGS3 IDRs from >200 plant species as a function of number of Trp residues present in the IDR. **(D)** Frequency of length distribution of the SGS3 IDRs included in the analysis. **(E)** Amino acid composition surrounding tryptophan residues in the analyzed set of IDRs.

### SDE3 knockout mutants have siRNA production defects different from those of AGO1 pocket mutants

Because knockout mutants in *SDE3*, unlike *SGS3,* have no defects in tasiRNA production (Vazquez *et al*., 2004), we reasoned that simple knockouts, rather than point mutants in Trp residues, would be sufficient for a first test of whether SDE3 is required for secondary siRNA production from miRNA target transcripts affected in AGO1 pocket mutants. Thus, we compared total small RNA profiles of C24 wild type and from CRISPR-Cas9-induced *sde3* mutant in this background (**Fig. EV5D-F**). This analysis revealed that only one of the miRNA-triggered secondary siRNA populations affected upon AGO1 pocket mutation showed reduced accumulation upon inactivation of *SDE3* (**Fig. 5A-D and Fig. EV5G**). Hence, SDE3 is unlikely to be the GW-interactor that explains the more widespread requirement for intact Trp-binding sites in AGO1 for miRNA-induced secondary siRNA generation, thus prompting us to design genetic tests for requirement of Trp-residues in the IDR of SGS3 for secondary siRNA production.

**Figure 5.**
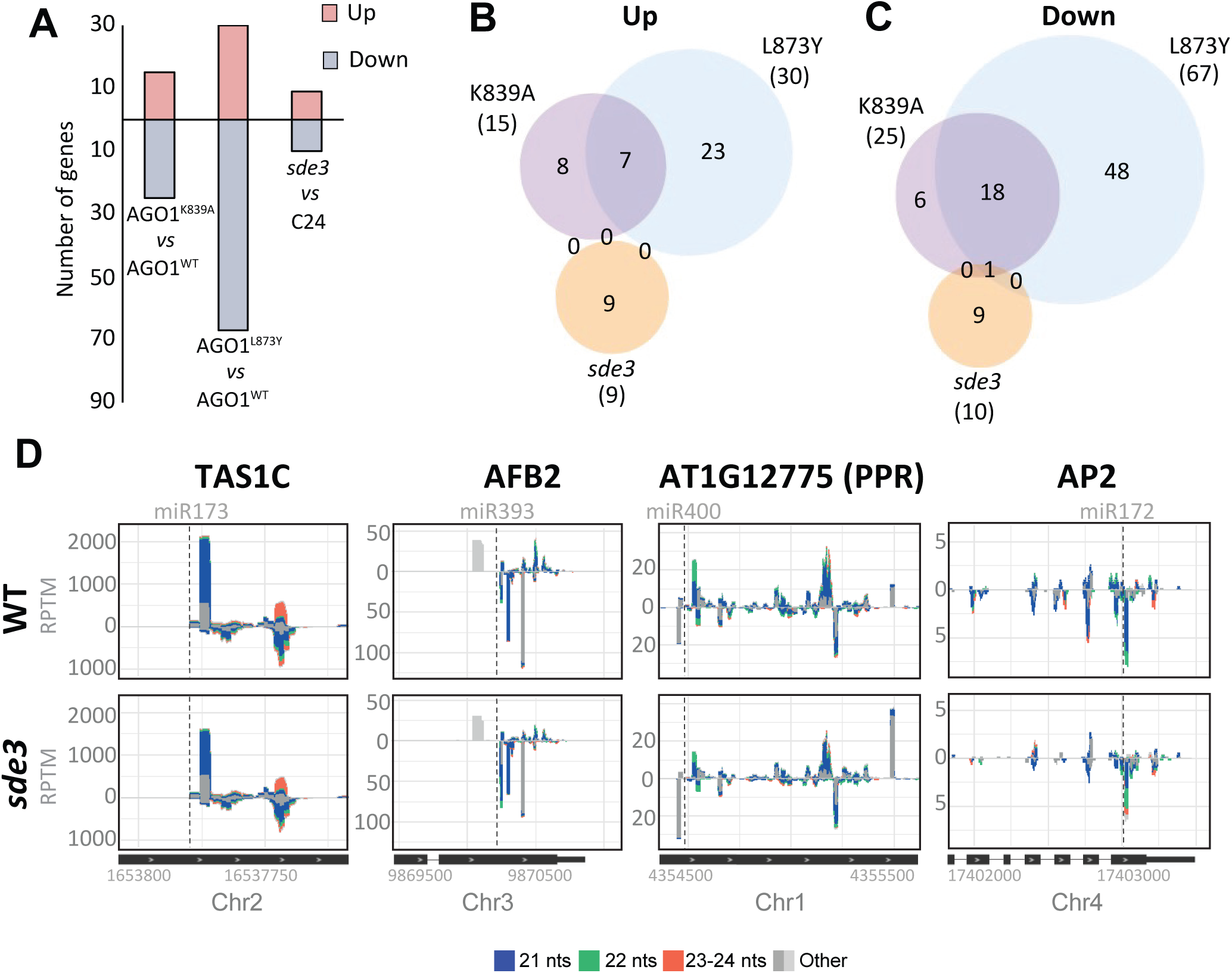
Secondary siRNA defects in *sde3* knockouts are distinct from those in AGO1 pocket mutants. **(A)** Bar chart of number of genes with differentially accumulated sRNAs in pocket 1 mutants (AGO1^K839A^ and AGO1^L873Y^) and the *sde3* mutant. **(B)-(C)** Venn diagrams illustrating the overlap of genes with upregulated (B) and downregulated **(C)** siRNA populations between the pocket 1 mutants (AGO1^K839A^ and AGO1^L873Y^) and the *sde3* mutant. **(D)** Examples of siRNA accumulation in Reads Per Ten Million (RPTM) for four representative genes in *sde3* and wild type samples. Gene models and genomic coordinates are shown below each plot (note: only partial gene models are displayed). miRNA target sites are highlighted with a dashed line. Small RNAs are color-coded by length: <21 nt (dark grey), 21 nt (blue), 22 nt (green), 23–24 nt (orange), and >24 nt (light grey).

### Mutation of IDR-located Trp residues in SGS3 reduces miRNA-induced secondary siRNA formation

Trp-dependent AGO interactions are typically tested genetically by Trp-Ala mutations in the interactor. Nonetheless, aromatic side chains in IDRs are also important for their propensity to form condensates (Martin *et al*, 2020), a property that may be important for SGS3-dependent siRNA amplification (Tan *et al*, 2023). We therefore engineered two quintuple point mutants in the five Trp residues in the N-terminal IDR of SGS3: one in which all Trp residues were changed to alanine (W44A/W63A/W74A/W137A/W139A, henceforth SGS3^5xA^) and one where they were changed to tyrosine (SGS3^5xY^, **Fig. 6**) to retain condensation propensity, but not the ability to bind to AGO1 pockets. Lines expressing similar levels of SGS3^WT^, SGS3^5xA^ and SGS3^5xY^ in the *sgs3-1* knockout background were further characterized. These lines showed normal levels of AGO1-dependent *TAS1/2* siRNAs and AGO7-dependent *TAS3* siRNAs, and complemented the leaf morphology phenotype of *sgs3-1* mutants caused by overexpression of the TAS3 targets *ARF3* and *ARF4* (Adenot *et al*, 2006; Fahlgren *et al*, 2006; Garcia *et al*, 2006; Hunter *et al*, 2006) (**Fig. 6A,C and Fig. EV6A-G**). Hence, the SGS3^5xA^ and SGS3^5xY^ mutant proteins do not abolish core SGS3 properties, probably including IDR-driven phase separation and localization to RDR6- and AGO7-containing siRNA bodies (Jouannet *et al*, 2012) that was previously suggested to be necessary for all SGS3 functions (Tan *et al*., 2023). Interestingly, however, the miR393-triggered *AFB2* secondary siRNAs affected in pocket mutants of AGO1 showed reduced abundance in the SGS3^5xA^- and SGS3^5xY^-expressing lines (**Fig. 6B-C and Fig. EV6E,D**). Furthermore, small RNA-seq showed that secondary siRNA production defects of the SGS3^5xA^ mutant across the entire transcriptome closely resembled those of pocket mutants in AGO1 (**Fig. 6D-I**). These observations support a model in which SGS3 binds to AGO1 via GW-pocket interactions that are necessary for many, but not all, cases of miRNA-induced secondary siRNA production. We note that the defective secondary siRNA production observed in the SGS3^5xY^ mutant supports a scenario where the Trp residues in SGS3 are important for specific protein-protein interactions rather than for merely dictating biophysical properties of the IDR.

**Figure 6.**
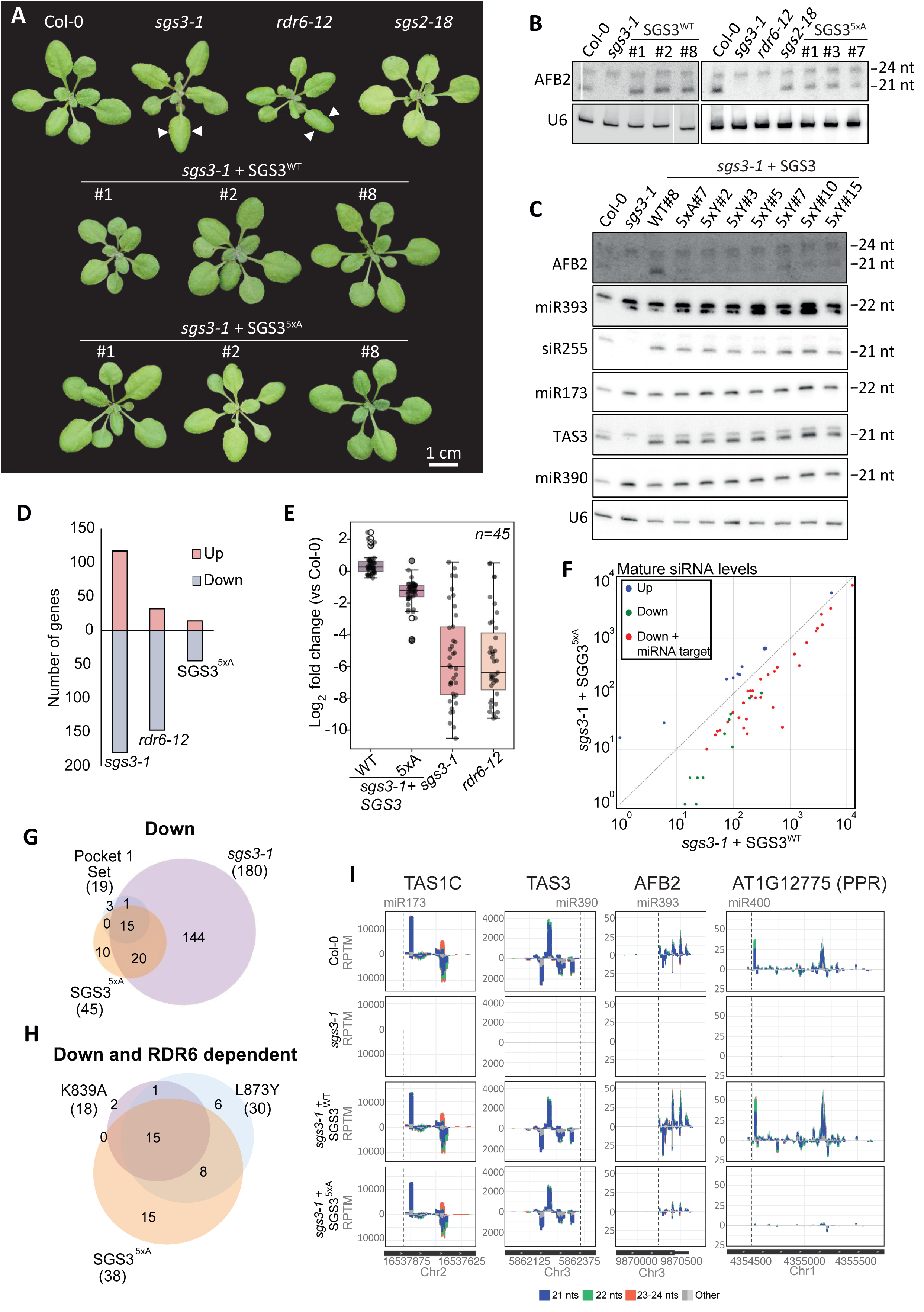
Mutation of Trp residues in the SGS3 IDR closely phenocopies secondary siRNA defects in AGO1 pocket mutants. **(A)** Pictures of 3.5-week-old Arabidopsis plants of Col-0 wild type (WT), *sgs3-1, rdr6-12*, *sgs2-18* and *sgs3-1* expressing the wild type genomic fragment of *SGS3* or the 5xA mutant in which the 5 Trp residues present in its IDR are mutated to Ala. Arrows indicate the downward-curved, pointy leaves characteristic of *rdr6-12*/*sgs3-1* mutants (Peragine *et al*, 2004). **(B)** RNA blot analysis of *AFB2* secondary siRNAs in inflorescences of the indicated genetic backgrounds. U6 served as a loading control. Lanes present on the original blot cropped out for presentation purposes are indicated by a dashed line. The entire blot is shown in Appendix Figure 2. **(C)** RNA blot analysis of secondary siRNAs and their corresponding trigger miRNAs in inflorescences of Col-0, *sgs3-1*, and *sgs3-1* lines expressing SGS3^WT^, SGS3^5xA^ and SGS3^5xY^. U6 serves as a loading control. **(D)** Histogram showing number of genes with differentially accumulated siRNAs in *sgs3-1*, *rdr6-12* and *sgs3-1* expressing SGS3^5xA^. *sgs3-1* and *rdr6-12* were compared to Col-0 plants while SGS3^5xA^ was compared to *sgs3-1* lines expressing transgenic SGS3^WT^. **(E)** Log_2_ fold change of siRNA accumulation in *sgs3-1*, *rdr6-12* or *sgs3-1* plants complemented with either the WT or 5xA SGS3 versions. The analysis includes siRNAs from the 45 genes identified by Deseq2 analysis in the comparison of SGS3^WT^ and SGS3^5xA^. **(F)** Scatter plot showing log_10_-transformed reads per ten millions (RPTM) values of siRNAs from the 45 individual genes analyzed in panel E, in *sgs3-1* lines complemented with either the SGS3^WT^ or SGS3^5xA^ versions. Blue indicates upregulation in SGS3^5xA^ compared to SGS3^WT^, green indicates downregulated siRNAs, and red indicated downregulation of siRNAs from miRNA target genes. **(G)** Venn diagram illustrating the overlap of genes with downregulated siRNA populations between (i) the pocket 1 gene set (AGO1^K839A^ ∩ AGO1^L873Y^ compared to AGO1^WT^), (ii) *sgs3-1* (compared to Col-0), and (iii) *sgs3-1* expressing SGS3^5xA^ (compared to *sgs3-1* expressing SGS3^WT^). **(H)** Venn diagram illustrating the overlap of genes with RDR6-dependent siRNAs downregulated in AGO1^K839A^, AGO1^L873Y^ and SGS3^5xA^. **(I)** Examples of secondary siRNA accumulation in Reads Per Ten Million (RPTM) for four genes in Col-0 wild type, *sgs3-1* and *sgs3-1* expressing either SGS3^WT^ or SGS3^5xA^. The four genes were selected to represent the pattern of affected and unaffected loci detected in AGO1 pocket mutants. *TAS1C* and *TAS3*, siRNA accumulation invariant upon AGO1 pocket mutation; *AFB2* and *AT1G12775*, siRNA accumulation reduced upon AGO1 pocket mutation. Gene models and genomic coordinates are shown below each plot (only partial gene models are displayed). miRNA target sites are highlighted with a dashed line. Small RNAs are color-coded by length: <21 nt (dark grey), 21 nt (blue), 22 nt (green), 23–24 nt (orange), and >24 nt (light grey).

We finally reasoned that if similarities in siRNA profiles between the AGO1 GW pocket mutants, SGS3^5xA^ and specific *rdr6* mutants that uncouple tasiRNA production from other siRNA amplifications could be found, they may reveal how the GW-mediated SGS3-AGO1 interaction facilitates RDR6 recruitment. We therefore examined siRNA profiles of *sgs2-18* mutants encoding the RDR6^P611L^ mutant protein that allows TAS3 siRNA production, yet is defective in transgene-derived siRNA amplification (Adenot *et al*., 2006). While *sgs2-18* mutants share some siRNA amplification defects with AGO1^K839^ and SGS3^5xA^ expressing lines (**Fig. EV4G-H and EV6H-I**), the overall effect on the siRNA transcriptome diverged substantially, suggesting that the GW pocket-mediated AGO1-SGS3 interaction contributes to RDR6 function through mechanisms distinct from those affected in *sgs2-18* mutants.

### The conserved negatively charged patch in the IDR of SGS3 is required for secondary siRNA formation

We next tested the functional importance of the negatively charged patch using a point mutant with four Asp/Glu→Lys/Arg mutations (SGS3^charge^) expressed in the *sgs3-1* knockout mutant background (**Fig. 7A**). Phenotypic analysis of lines expressing SGS3^charge^ showed that similar to SGS3^5xA^, they complemented leaf morphology phenotypes of *sgs3-1* knockout mutants (**Fig. 7B**). Importantly, the accumulation of miR393-induced *AFB2* secondary siRNAs was nearly completely abrogated in the SGS3^charge^ mutants, with little or no effect on *TAS1C* and *TAS3* siRNAs (**Fig. 7C**). Thus, mutation of (i) the Trp binding pockets in AGO1, (ii) the surrounding positively charged surface, (iii) the Trp residues in the IDR of SGS3 or (iv) its conserved negatively charged patch produces very similar defects in secondary siRNA production, suggesting that an SGS3-GW:AGO1 binding facilitated by polyelectrolyte interactions is required for secondary siRNA formation.

**Figure 7.**
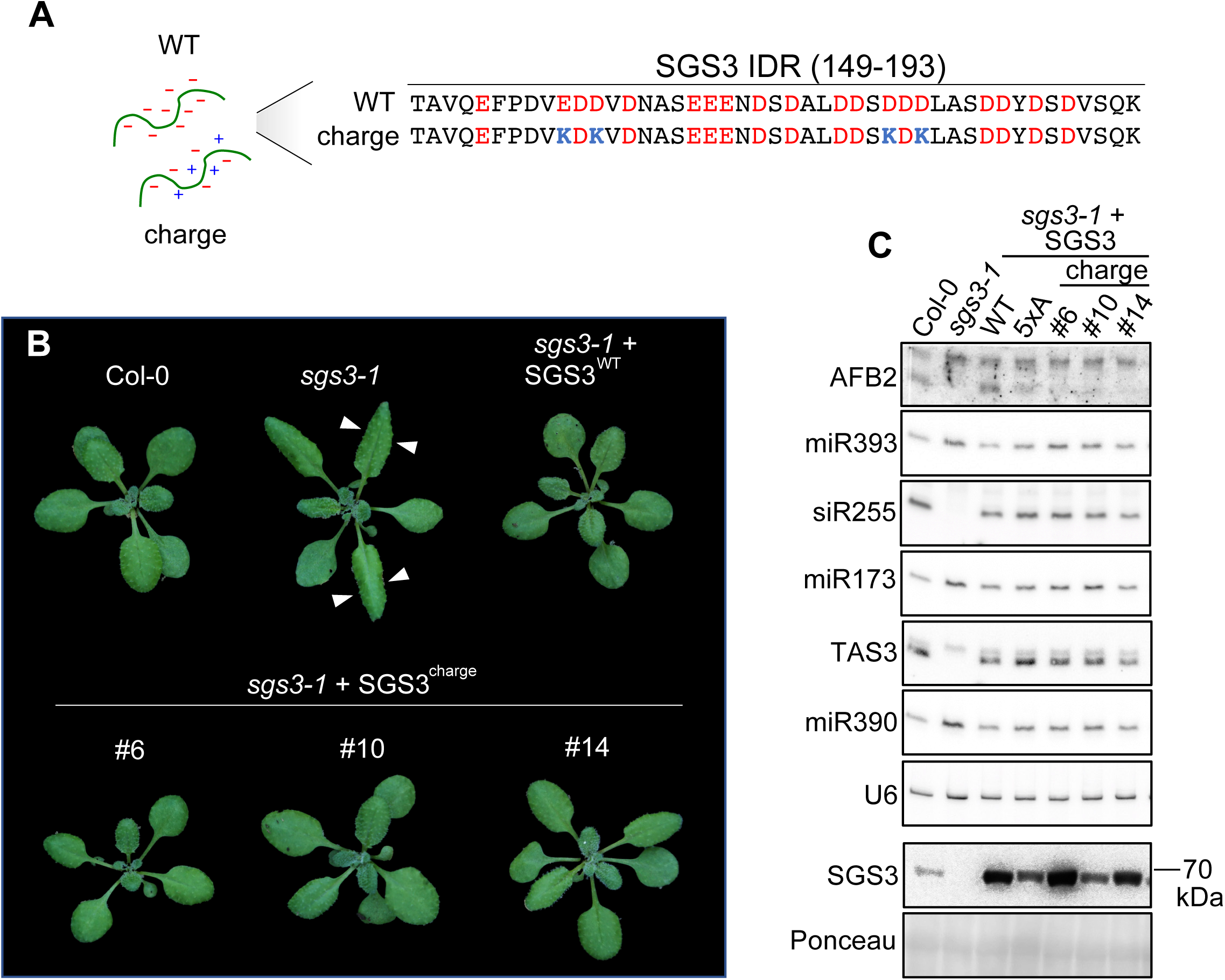
Disruption of the negatively charged patch in the SGS3 IDR causes secondary siRNA defects similar to those in AGO1 pocket mutants. **(A)** Amino acid sequence of a C-terminal part of the SGS3 IDR (residues 149-193) highlighting negatively charged residues (red) and those mutated to lysine in the charge mutant (blue). **(B)** Representative images of 3.5-week-old Arabidopsis plants of the indicated genetic backgrounds. Three independent transgenic lines of *sgs3-1* expressing SGS3^charge^ are shown. Arrows indicate the downward-curved, pointy leaves of *sgs3-1* (Peragine *et al*., 2004). **(C)** RNA blot analysis of secondary siRNAs and their corresponding trigger miRNAs in inflorescences of Col-0, *sgs3-1*, and *sgs3-1* transgeneic lines expressing *SGS3^WT^*, *SGS3^5xA^* or *SGS3^charge^*. U6 serves as a loading control. SGS3 protein levels were analyzed by immunoblot from the same samples. Ponceau staining was used as a loading control.

### Properties of TAS precursors, not of the miR173 trigger, underlie the independence of GW-pockets for tasiRNA production

The model of a GW:pocket-mediated SGS3-AGO1 interaction required for secondary siRNA production has one obvious flaw. It does not explain why mutations in GW pockets and Trp residues in IDR^SGS3^ have no impact on *TAS1ABC* and *TAS2* tasiRNA production triggered by miR173. In principle, the explanation could be found either in special properties of the miR173 trigger, in the *TAS1ABC/TAS2* precursor transcripts, or in both. To distinguish between these possibilities, we used a synthetic *TAS1C*-based tasiRNA (syn-tasiRNA) system in which the 2^nd^ siRNA generated from the miR173 cleavage site is replaced by an siRNA targeting the *CHLORINA42* (*CH42*) mRNA encoding a Mg^II^-chelatase subunit (Cisneros *et al*, 2025; Felippes & Weigel, 2009), known as *SULPHUR* (*SUL*) in *Nicotiana tabacum*. This variant of the *SUL* silencing system (Himber *et al*., 2003), syn-tasiSUL, enables detection of secondary siRNA production by visual inspection of leaf yellowing (**Fig. 8A-B**), and is appropriate for tests involving AGO1 pocket mutants because they do not have different levels of miR173 compared to wild type (**Fig. 8C**). We first showed that a *MIR173* deletion mutant fully suppressed *CH42* silencing (**Fig.8D-E**) indicating that the syn-tasiSUL system specifically monitors miR173-induced secondary siRNA activity, and that potential contributions to *CH42* silencing from co-suppression of the syn-tasiSUL construct are negligible. We then introduced two variants of the syn-tasiSUL system into our set of AGO1 pocket mutants: one with the full *TAS1C*-based syn-tasiSUL precursor and a minimal version containing only the miR173 target site, an 11-nt linker and the SUL siRNA (Cisneros *et al*., 2025). In the presence of wild type AGO1, either FLAG-AGO1^WT^ or endogenous AGO1, both versions induced efficient *CH42* silencing in primary transformants (**Fig.8D-E**). Importantly, in the AGO1 pocket mutants, the frequency of *CH42* silencing was clearly reduced only for the minimal version (**Fig. 8D-E**). Hence, the ability of miR173/*TAS* precursor targeting to sustain secondary siRNA production in the absence of GW-mediated AGO1-SGS3 interaction is explained by properties in the *TAS* precursors other than the miR173 target site.

**Figure 8.**
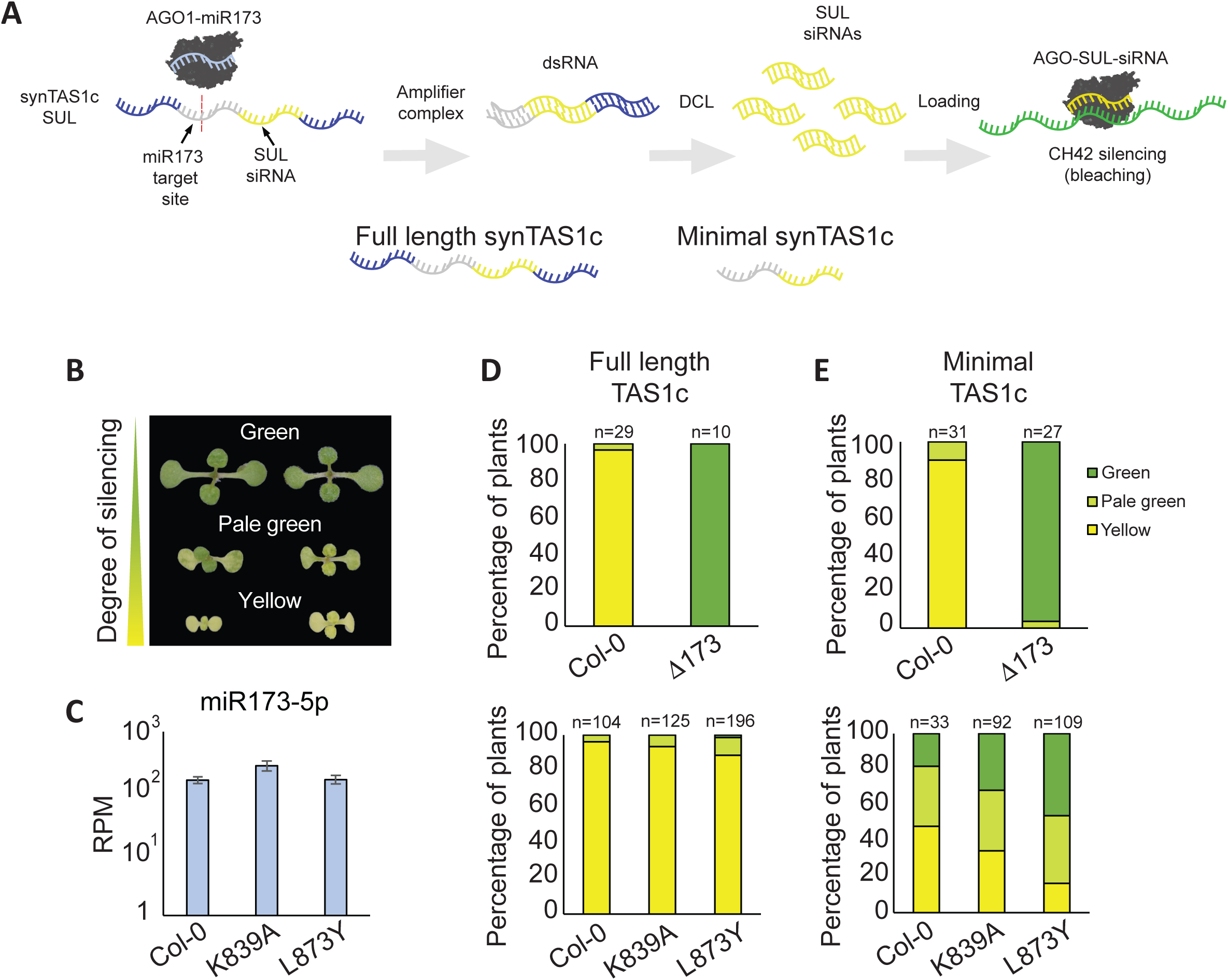
*TAS1C* target RNA properties underlie the AGO1 pocket independence of its secondary siRNAs. **(A)** Schematic of *CH42* silencing by syn-*TAS1C*-derived siRNAs (Cisneros *et al*., 2025). A synthetic *TAS1C* precursor (syn-*TAS1C*-SUL) was engineered to include a 21-nt fragment of the *CH42* mRNA sequence. Upon recognition and cleavage by the AGO1–miR173 complex, the 3′ cleavage fragment is converted into double-stranded RNA via SGS3-SDE5-RDR6 and subsequently processed into siRNAs by DCL4. The resulting *SUL*-specific siRNA is incorporated into AGO proteins and mediates post-transcriptional silencing of *CH42*. Two synthetic precursors were used in this study: (i) a full-length syn-*TAS1C* construct mimicking the endogenous TAS1C architecture, and (ii) a minimal syn-*TAS1C* version containing only the miR173 target site, a linker and the *CH42* fragment. **(B)** Representative phenotypes and scoring criteria used to assess CH42 silencing in first transgenic generation (T1) plants transformed with the full-length *TAS1C* precursor. Silencing categories were defined as: Green (no silencing), Pale green (intermediate silencing), and Yellow (strong silencing). **(C)** miR173 levels in Col-0 (WT) and *ago1-3* lines expressing either FLAG-AGO1^K839A^ or FLAG-AGO1^L873Y^ (pocket 1 mutants). Values in RPMs (Read Per Million). miR173 reads were retrieved from libraries generated in this study. **(D)** Quantification of phenotypic categories observed in T1 populations transformed with the full-length *TAS1C* precursor in Col-0, *mir173* mutant (Δ173), and *ago1-3* lines expressing FLAG-AGO1^K839A^ or FLAG-AGO1^L873Y^. The number of analyzed T1 individuals is indicated above each bar (n). **(E)** Quantification of phenotypic categories observed in T1 populations transformed with the minimal *TAS1C* precursor in the same genotypes as in (D). The number of T1 individuals scored is indicated above each bar.

### The SGS3^GW^-dependent amplification mechanism is required for amplified RNAi

We finally tested whether the SGS3^GW^-dependent mechanism is implicated in amplified RNAi against foreign genetic elements, the likely ancestral role of RDR6-mediated siRNA amplification. To do so, we employed the well-established β-glucuronidase (GUS) transgene silencing system *L1* in the *sgs3-1* knockout background (Mourrain *et al*., 2000), and used crossing to introduce either a endogenous wild type *SGS3* allele, or *SGS3^5xA^* or *SGS3^Charge^* transgenes expressed in the *sgs3-1* background. Because *L1* silencing is sensitive to SGS3 dosage as seen by the different kinetics of *L1* silencing in *SGS3/SGS3* homozygous wild type and *SGS3/sgs3-1* mutants heterozygous for the *sgs3-1* knockout allele (**Fig. EV7A**), we chose *SGS3^5xA^* and *SGS3^Charge^* transgenic lines with expression levels similar or moderately higher than the level of SGS3 in Col-0 wild type. In both cases, none of the SGS3^5xA^-or SGS3^Charge^-expressing plants reached full GUS silencing at any point during the 63-day time course analysis, contrary to SGS3 wild type for which GUS expression in every plant was eventually completely silenced (**Fig. 9A**). Consistent with this clear silencing phenotype, GUS siRNA levels were high in plants expressing SGS3 wild type, but undetectable in the SGS3^5xA^-or SGS3^Charge^-expressing plants (**Fig. 9B**). We also analysed endogenous tasiRNA populations to verify that SGS3^5xA^ and SGS3^Charge^ sustained the ability to produce tasiRNAs in rosette leaves, as previously seen in flowers. For SGS3^5xA^, we observed wild type levels of *TAS1* and *TAS3* siRNAs as expected (**Fig. 9B**). Unexpectedly, SGS3^Charge^ conferred a partial reduction in *TAS1* and *TAS3* siRNAs in rosettes (**Fig. 9B**), suggesting that the negatively charged patch serves functions in addition to facilitation of the interaction with AGO1 pockets and that this is detected in leaves that have lower SGS3 expression than flowers. Indeed, when additional mutations in the negatively charged patch were introduced, we also observed reduced *TAS1* and *TAS3* siRNA levels in flowers (**Fig. EV8A-C**). We conclude that the SGS3^GW^-mediated mechanism, presumably relying on direct interaction with the Trp-binding pockets of AGO1, is required for fully functional siRNA amplification in RNAi.

**Figure 9.**
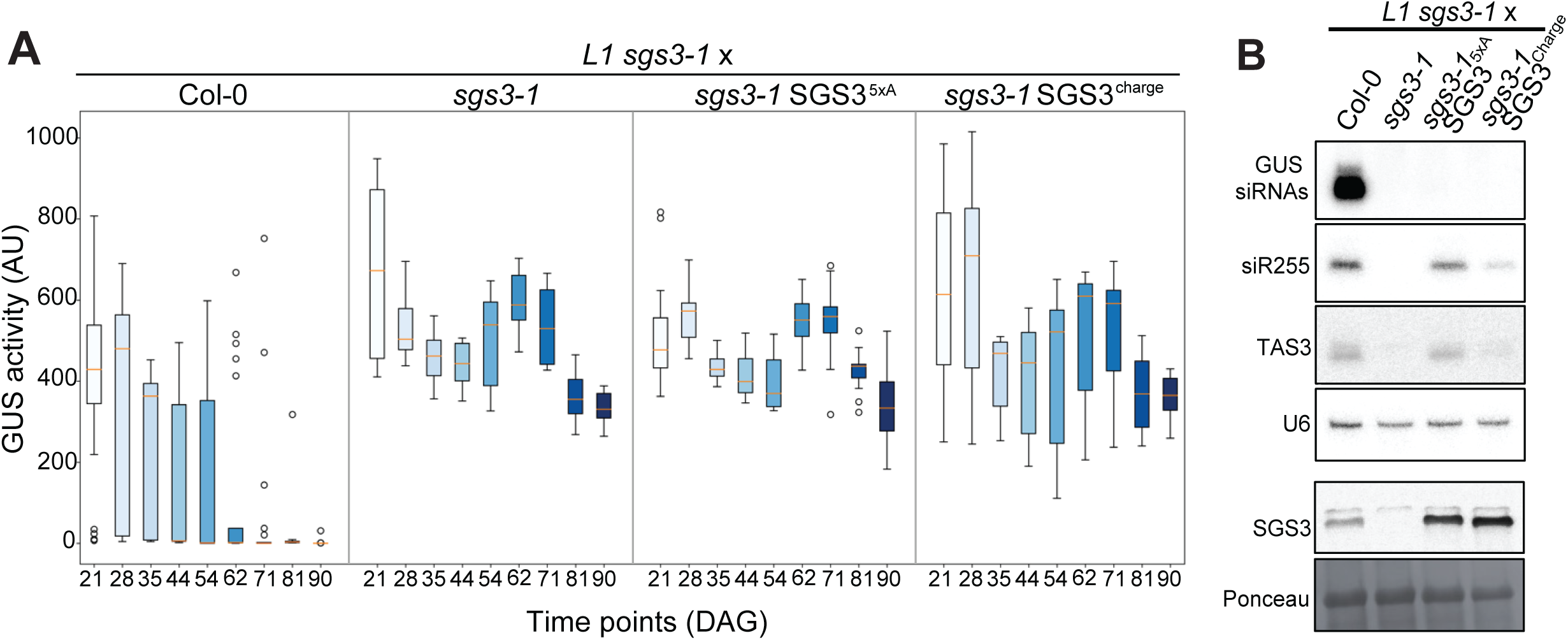
SGS3^GW^-dependent siRNA amplification is essential for transgene silencing. **(A)** Kinetics of post-transcriptional gene silencing of GUS (*L1* line) measured as GUS activity (arbitrary fluorescence units). Lines inside boxes indicate the median (Q2) GUS activity, and the length of boxes indicate the interquartile range (IQR = Q3-Q1). The whiskers extend outwards to the smallest and the largest data points within 1.5 x IQR from Q1 and Q3, respectively. Samples were collected from F1 populations of crosses between *L1*/*sgs3-1* and Col-0, *sgs3-1*, *sgs3-1*/SGS3^5xA^ and *sgs3-1*/SGS3^charge^. Samples were collected at the indicated days after germination. **(B)** RNA blot analysis of GUS-derived siRNAs and endogenous secondary siRNAs (siR255 and TAS3) in leaves of plants of the genotypes analysed in (A). U6 served as a loading control. SGS3 protein levels were analyzed by immunoblot from the same samples. Ponceau staining was used as a loading control.

## DISCUSSION

### siRNA amplification via RISC-mediated amplifier complex assembly, not via mere generation of cleavage fragments

The evidence that RISC itself rather than cleavage fragments released after RISC-catalyzed slicing triggers siRNA amplification is now solid. For example, this RISC trigger model (Branscheid *et al*., 2015) is supported by the observation that miR173 can trigger tasiRNA production in the absence of catalytic activity of AGO1 (Arribas-Hernandez *et al*., 2016b), and we show here that siRNA production triggered by other miRNAs depends strongly, in some cases fully, on intact GW-binding pockets, even if slicer activity is intact. Thus, small RNA-guided cleavage activity and the ability to trigger secondary siRNA production can be fully genetically uncoupled. It follows that the originally suggested molecular function of SGS3, stabilization of RISC cleavage fragments for use as RDR6 templates (Yoshikawa *et al*., 2005), is incomplete, or perhaps even misleading. The observations that SGS3 is required for secondary siRNA production in the absence of slicing (Arribas-Hernandez *et al*., 2016b; de Felippes *et al*, 2017), and that the AGO1:RDR6 association requires SGS3 (Sakurai *et al*., 2021; Yoshikawa *et al*., 2021) argue that the key function of SGS3 is to form an RDR6-containing amplifier complex on RISC target RNA. In this model, the loss of RISC cleavage fragments in *sgs3*, but not in *rdr6*, knockout mutants may be a secondary effect of loss of the factor that keeps RISC on the cleaved target RNA for amplification: without SGS3, RISC dissociates from the target RNA allowing concomitant cleavage fragment degradation, for instance by SKI2-3-8-exosome and XRN4-mediated exonucleolysis (Branscheid *et al*., 2015; Souret *et al*, 2004; Vigh *et al*, 2022).

### Is the SGS3^GW^-AGO1 interaction direct?

The mutational evidence provided here, together with the previously reported evidence for physical association between SGS3 and AGO1 (Sakurai *et al*., 2021; Yoshikawa *et al*., 2021; Yoshikawa *et al*., 2013), supports a model in which the GW motifs in IDR^SGS3^ directly interact with the Trp-binding pockets of AGO1. Clearly, this model is also consistent with the structurally well-established role of Trp-binding pockets in eukaryotic Ago proteins (Elkayam *et al*., 2017; Pfaff *et al*., 2013; Schirle & MacRae, 2012; Sheu-Gruttadauria & MacRae, 2018). We favor this model because it readily explains (i) the remarkably similar effects on secondary siRNA production in AGO1 pocket and SGS3^5xA^ mutants, (ii) the requirement for positively charged residues around the pockets in AGO1 and for negatively charged residues in the IDR^SGS3^, (iii) why miR393-induced secondary siRNA production is defective when Trp residues in IDR^SGS3^ are replaced by Tyr residues that would also promote macromolecular condensation. Nonetheless, our attempts at directly proving such an SGS3-AGO1 interaction with different fragments of IDR^SGS3^ were unsuccessful, because in most preparations, the IDR^SGS3^ fragments tended to undergo condensation and indiscriminate association with many proteins. Hence, with the current evidence at hand, we consider the results to be best explained by a direct SGS3^GW^-AGO1 interaction but cannot formally exclude the existence of unidentified helper proteins in the physical association between SGS3 and AGO1. Nonetheless, the remainder of the discussion treats implications of our results based on the direct SGS3^GW^-AGO1 interaction model.

### Two distinct modes of SGS3 recruitment to AGO1

The core biochemical activity of SGS3 that allows association with target-bound RISC is its affinity for dsRNA with long 5’-overhangs (Fukunaga & Doudna, 2009), consistent with its binding to the 3’-end of AGO-bound small RNA in the presence of a complementary target RNA (Iwakawa *et al*, 2021). We propose that this activity is insufficient for stable binding of SGS3 to target-bound RISC and that at least two additional mechanisms can ensure the stable binding required for RDR6 recruitment: one is GW-mediated SGS3:AGO1 interaction, but our analysis of tasiRNA production clearly shows that this cannot be the only possible mechanism. The *TAS* precursor transcripts contain a short open reading frame critical for secondary siRNA formation immediately upstream of the miR173 binding site (Yoshikawa *et al*, 2016; Zhang *et al*, 2012). Similarly, TAS3 tasiRNAs induced via the AGO7-miR390 complex require a short open reading frame with a strong ribosome stall site juxtaposed to the miR390 target site, and this juxtaposition is required for SGS3:AGO7 interaction (Sakurai *et al*., 2021). Such strong ribosome stall sites are not present in the miRNA targets from which secondary siRNA formation requires the GW-mediated SGS3:AGO1 interaction. Hence, we propose that stable SGS3:AGO1 interaction can be achieved either by GW-mediated binding to the hydrophobic pockets in AGO1, or by a currently ill-defined mechanism that relies on a stalled ribosome juxtaposed to RISC. Clearly, it is a key objective of future work to identify this second AGO1:SGS3 interaction site and to analyse whether abolishment of AGO1:SGS3 interaction altogether leads to complete loss of RDR6-dependent siRNA amplification via AGO1-bound small RNAs, a key prediction of the RISC trigger model for siRNA amplification.

### The GW-pocket interaction as a SLiM-driven combinatorial protein-protein interaction

Our study reveals that at least two structural AGO1 elements are required for the function of the GW-binding pockets in secondary siRNA production: First, a positively charged surface around the pockets likely engages in polyelectrolyte interactions with the negative patch in IDR^SGS3^, explaining why it is as important as the pockets for secondary siRNA production. Second, the β-finger is also strictly required, although our study did not define the molecular basis of this requirement. The sequence of the β-finger is highly conserved across AGO1 orthologues (Xiao *et al*., 2023), consistent with a function in direct binding to a conserved interactor such as SGS3. Binding to another component in siRNA amplification, perhaps SDE5, or a more indirect effect on pocket function cannot be excluded at this point, however. Notwithstanding the currently undefined molecular function of the β-finger, the finding of accessory sites on AGO1 required for pocket function suggests that the GW-AGO interaction follows the logic of many other interactions between short linear motifs (SLiMs) in IDRs and globular domains: each individual interaction is weak, and stable association requires a combination of several such weak interactions. In the case of GW-AGO interactions, this combinatorial mode may simply be restricted to several minimal GW SLiMs binding to the pockets on the same or closely spaced AGO proteins, as has been shown for TNRC6/HsAgo1/2 (Elkayam *et al*., 2017; Sheu-Gruttadauria & MacRae, 2018). However, the AGO1:SGS3 association studied here provides a clear example that other molecular interactions may determine whether a possible GW-pocket interaction leads to macromolecular association. This is clearly of critical importance in organisms like higher plants where different AGO proteins use GW-based interactions with different partner proteins to achieve completely different biological outputs.

### Relation to other proposed GW interactors of AGO1 in plants

Previous studies have proposed three different proteins as endogenous GW-interactors of AGO1: the nuclear protein SUO (Yang *et al*, 2012), the cytochrome P450 enzyme ROT3 (Wang *et al*, 2021) required for brassinosteroid biosynthesis (Kim *et al*, 1998), and the RNA helicase SDE3 (Garcia *et al*., 2012), required for some cases of siRNA amplification (Dalmay *et al*., 2001; Himber *et al*., 2003; Jauvion *et al*., 2010). Physical association between SUO and AGO1 has not been demonstrated, and the Trp residues in SUO are located in folded regions not expected to be available for interaction, suggesting that SUO does not bind to AGO1 via GW-pocket interactions. Co-immunoprecipitation of ROT3 and SDE3 with AGO1 requires GW dipeptides in predicted IDRs, but in both cases, the effect of mutating the hydrophobic pockets in AGO1 was not tested (Garcia *et al*., 2012; Wang *et al*., 2021). Because aromatic residues in IDRs are key drivers of biomolecular condensation (Mittag & Pappu, 2022), Trp-dependent co-immunoprecipitation with AGO1 from total lysates alone is not strong evidence for a direct GW-pocket interaction. In particular, for ROT3, the Trp-dependent AGO1 association is unlikely to involve the hydrophobic pockets in AGO1, because it was reported to influence the miR398-guided translational repression of CSD2 (Wang *et al*., 2021), a process that we found to be unaltered in pocket 1 mutants of AGO1. The previous evidence for direct GW-mediated SDE3:AGO1 interaction (Garcia *et al*., 2012) combined with our observation that a GW-containing SDE3 peptide requires an intact pocket 1 for full AGO1 binding in pull-down assays supports the notion that SDE3 is a *bona fide* GW interactor. This raises an immediate problem: how can two proteins implicated in siRNA amplification be competing for the same binding site in AGO1? One possibility may be that SGS3 and SDE3 do not, in fact, compete for binding even if they use the same binding site in AGO1. siRNA amplification may require physical proximity of RISC loaded with the trigger small RNA and of an RNA-free AGO1 to accept amplified siRNA. Since GW pockets are not expected to be affected by small RNA binding, it is possible that SGS3 stimulates siRNA amplification by interaction with the trigger RISC while SDE3 does so by bringing RNA-free AGO1 to the site of amplification. If so, SDE3 may use combinatorial AGO1 interaction relying on GW pockets and one or more elements uniquely accessible in the RNA-free state, a possibility that is now testable with the recent identification of a structural switch in eukaryotic AGO proteins available for interaction only in the RNA-free state (Bressendorff *et al*, 2025).

### Similar biochemistry, divergent biology: cooperativity through AGO GW pockets and partners

Plant AGO1 is the orthologue of metazoan miRNA-associated Ago proteins, including human Ago2. Both are major miRNA effectors and both proteins have conserved GW binding pockets. Yet, these highly conserved binding sites have acquired divergent functions: Trp-dependent interaction with the metazoan-specific GW182/TNRC6-family for miRNA-guided target repression in the case of metazoan miRNA-associated Ago proteins, and Trp-dependent interaction with plant-specific SGS3 for siRNA amplification in plants. Intriguingly, the underlying molecular logic of these interactions may have similarities linked to cooperative RISC action. In metazoans, miRNA target sites present in the same mRNA have long been known to confer mRNA repression cooperatively (Doench & Sharp, 2004; Vella *et al*, 2004), and recent biochemical analyses show that GW-mediated TNRC6-AGO interaction underlies this effect as cooperative RISC binding to target RNA can be recapitulated *in vitro* in the presence of the GW-containing AGO-interacting part of TNRC6 (Briskin *et al*, 2020). Similarly, 21-nt miRNA target sites in plants show cooperative effects on secondary siRNA formation (Axtell *et al*., 2006; Howell *et al*., 2007). Two lines of evidence suggest that the GW-mediated SGS3:AGO1 interaction may underlie this cooperative effect.

First, we show here that mutation of the pockets 1 and 3 in AGO1 and of Trp residues in IDR^SGS3^ abolishes this cooperativity. Second, SGS3 acts as a dimer or higher-order multimer (Elmayan *et al*, 2009), and IDR^SGS3^ has several possible AGO1-interacting Trp residues, suggesting that a single SGS3 dimer/multimer could interact with several RISCs on the same target RNA, thereby causing cooperative binding. Hence, a basic function of GW binding pockets may be to enable cooperative RISC action, although this is now used to different effects in plants and metazoans. We note, however, that SGS3 is unlikely to be the only GW-interactor of AGO1 in plants, and that TNRC6/GW182-independent metazoan miRISC effects have been reported (Bukhari *et al*, 2016; Jannot *et al*, 2016; Matsuura-Suzuki *et al*, 2024; Wu *et al*, 2013). Thus, the interesting possibility that an ancestral interactor of the highly conserved GW binding pockets common to both classes of Ago proteins remains to be discovered should not be completely excluded, although it is also possible that such an ancestral interactor was independently lost and replaced by others during the course of evolution.

## EXTENDED VIEW FIGURE LEGENDS

**Figure EV1.**
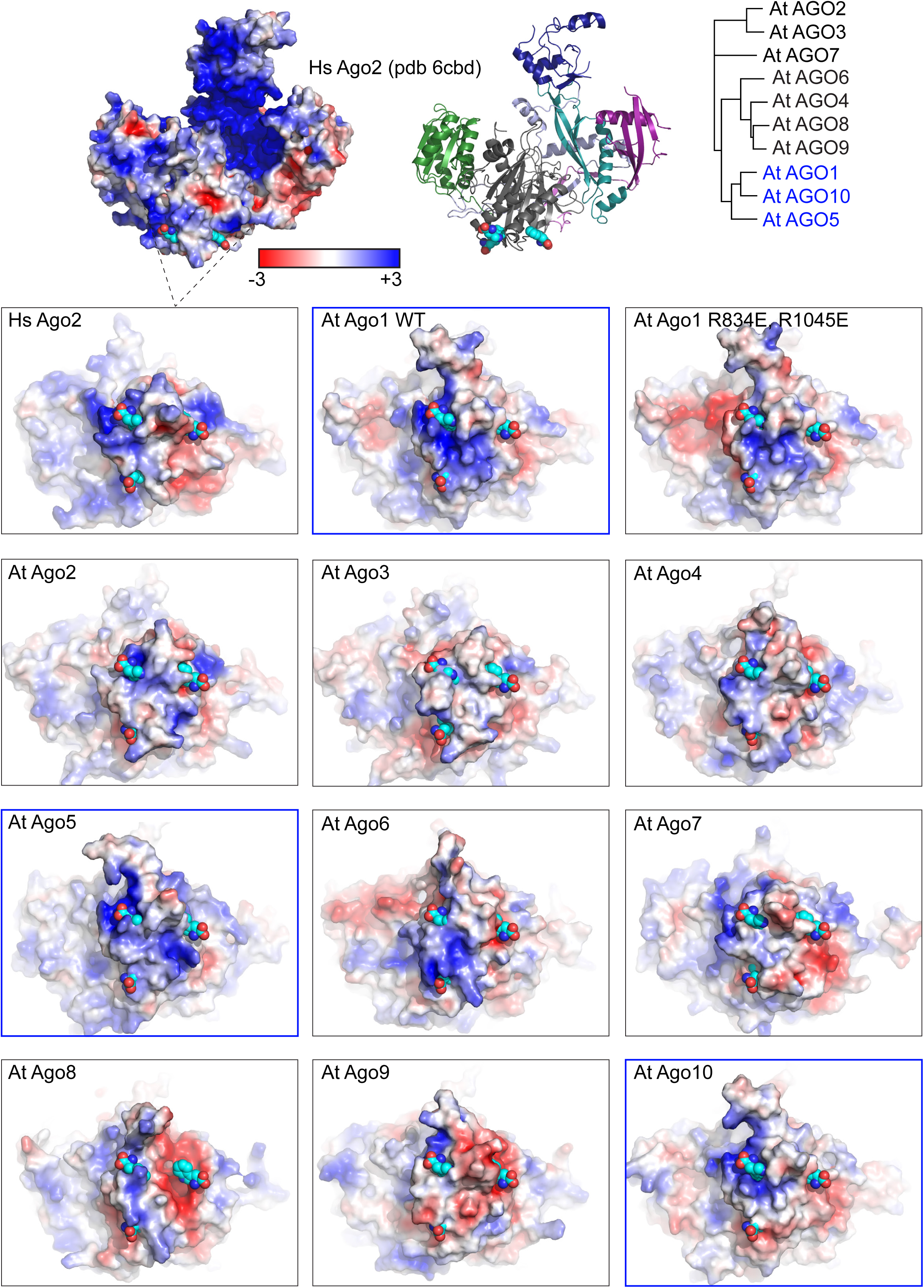
Structural overview and charge distribution of AtAGO pocket regions and surrounding elements. **(A)** Structural overview of the pocket region in HsAGO2, AtAGO1, AtAGO1^RR-EE^, and AtAGO2-10 proteins. Structural models were obtained from AlphaFold3 predictions. Amino acids are colored on a red (negative) to blue (positive) scale according to their charge. Tryptophan residues accommodated within the pockets are shown in cyan. A phylogenetic tree of AtAGO1 to AtAGO10 is shown in the right upper panel.

**Figure EV2.**
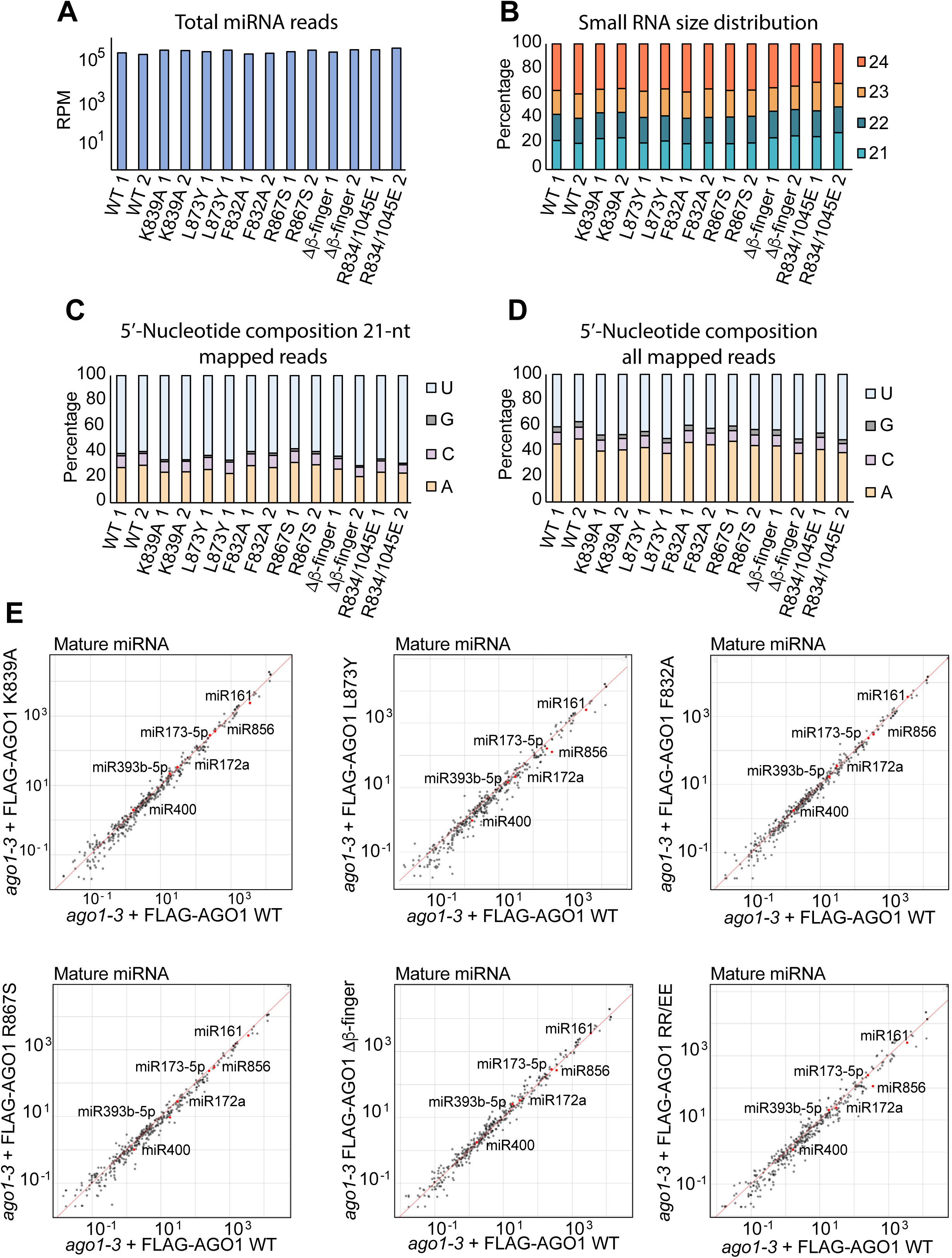
Mutations in AtAGO1 pockets or surrounding structural elements do not affect miRNA profiles. **(A)** Total miRNA reads in Reads Per Million (RPMs) in *ago1-3* plants expressing either FLAG-AGO1^WT^ or the indicated FLAG-AGO1 mutants with amino acid changes in the pocket region. **(B)** Small RNA size distribution (percentage) of 21 to 24-nt species of all mapped reads for the genotypes analyzed in (A). **(C)** 5’-nucleotide composition of all the 21-nt mapped small RNA reads in plants of the genotypes analyzed in (A). **(D)** 5’-nucleotide composition of all mapped small RNA reads in the same genotypes analyzed in (C). **(E)** Scatter plots showing log₁₀-transformed reads per million (RPM) values of individual miRNAs detected in *ago1-3* plants complemented with either FLAG-AGO1^WT^ or FLAG-AGO1 pocket-mutated versions. miRNAs of particular relevance to this study are highlighted in red. The red diagonal line represents equal abundance in both genotypes.

**Figure EV3.**
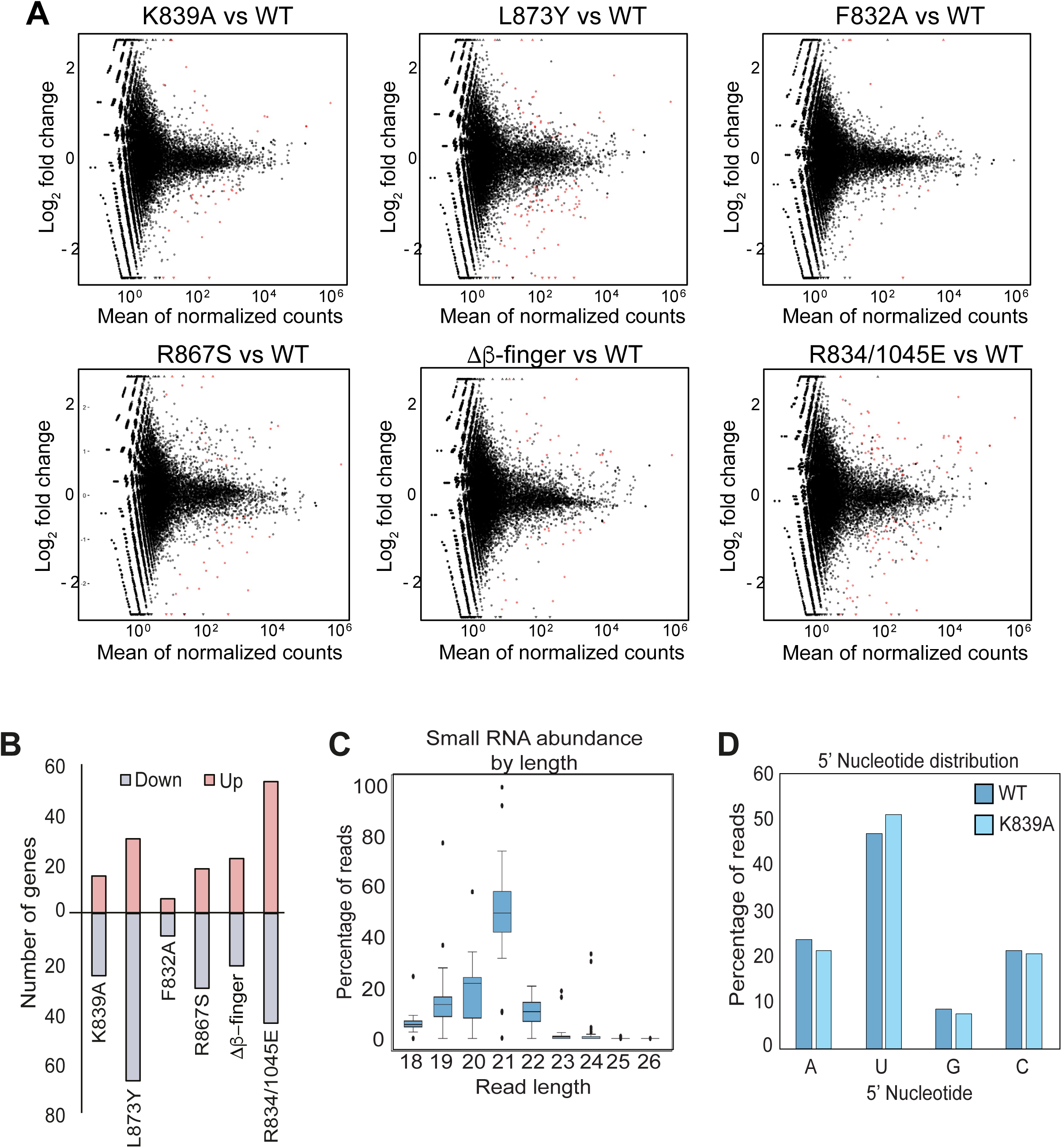
Mutations in AtAGO1 pocket regions or surrounding structural elements affect siRNA production at multiple loci. **(A)** MA plot showing differential accumulation of small RNAs from genes (red dots) in *ago1-3* plants expressing either FLAG-AGO1^WT^ or pocket-mutated FLAG-AGO1 variants. Statistical analyses were performed by Deseq2. **(B)** Bar chart of number of genes with differentially accumulated sRNAs in in *ago1-3* plants complemented with FLAG-AGO1 variants. Statistical analysis was performed by Deseq2 using FLAG-AGO1^WT^ plants as a control. **(C)** Size distribution of small RNAs mapping to the downregulated pocket 1 gene set defined in Figure 3D. Reads from the 19 genes were retrieved and the percentage of each size class was calculated. **(D)** 5′-nucleotide composition of the small RNAs analyzed in (C).

**Figure EV4.**
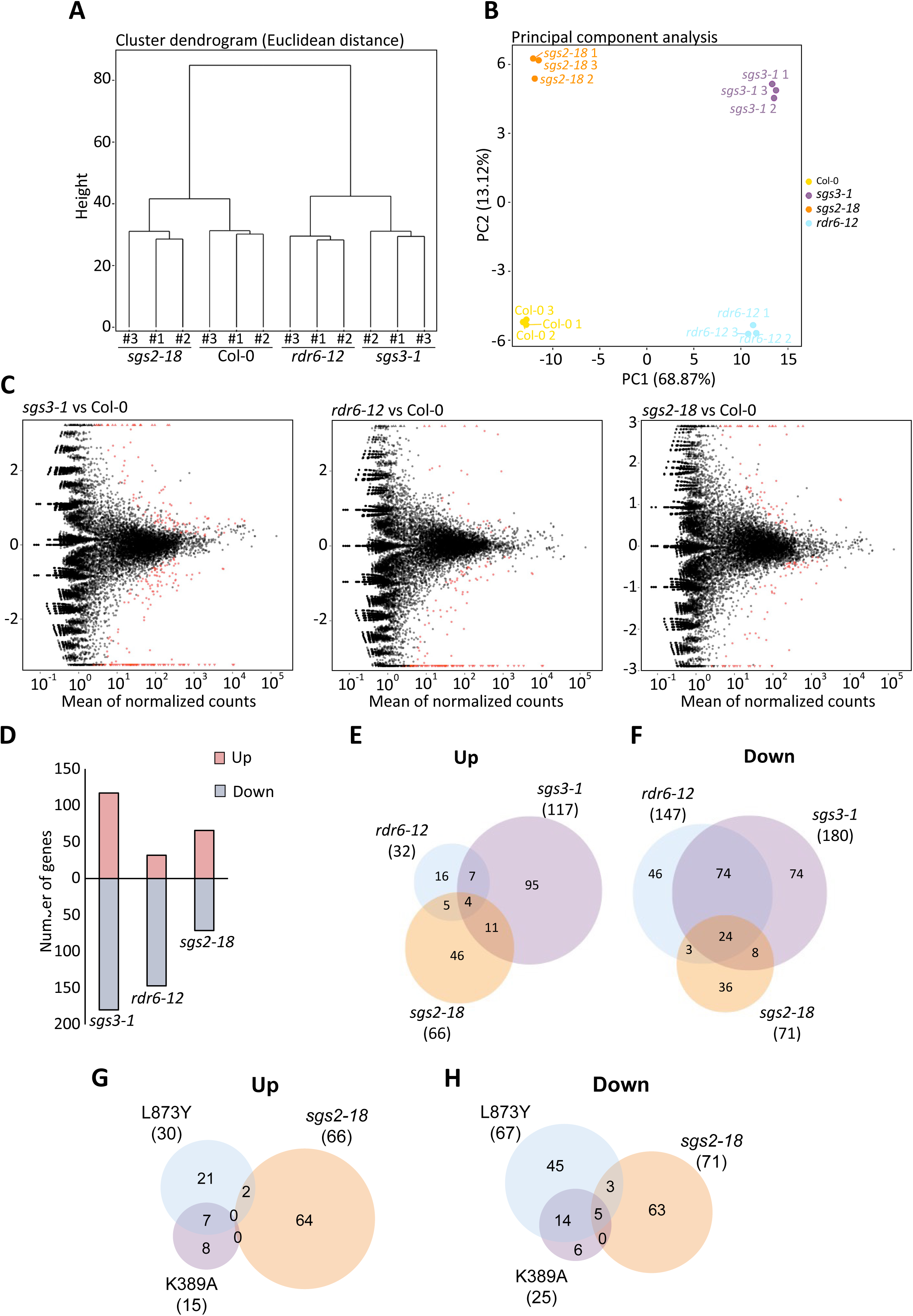
Small RNA-seq analysis of mutants impaired in siRNA amplification. **(A)** Hierarchical clustering of small RNA-seq data based on Euclidean distances between biological replicates of Col-0 (Wild-type), *sgs3-1*, *rdr6-12,* and *sgs2-18*. **(B)** Principal component analysis of three biological replicates analyzed in (A). **(C)** MA plot showing differential accumulation of small RNAs from genes (red dots) in *sgs3-1, rdr6-12 and sgs2-18* compared to Col-0 wild type. **(D)** Bar chart of number of genes with differentially accumulated small RNAs in *sgs3-1, rdr6-12* and *sgs2-18* relative to Col-0 wild type. **(E)** Venn diagrams illustrating the overlap of up and **(F)** downregulated genes between *sgs3-1*, *rdr6-12, and sgs2-18*. Gene sets were derived from transcriptomic comparisons relative to Col-0 Wild-type. **(G)-(H)** Venn diagrams illustrating the overlap of genes giving rise up-(G) and downregulated (H) siRNAs between *sgs2-18*, or *ago1-3* plants expressing either FLAG-AGO1^K839A^ or FLAG-AGO1^L873Y^ versions. Gene sets were derived from transcriptomic comparisons relative to Col-0 Wild-type for *sgs2-18*, and FLAG-AGO1^WT^ for K839A and L873Y.

**Figure EV5.**
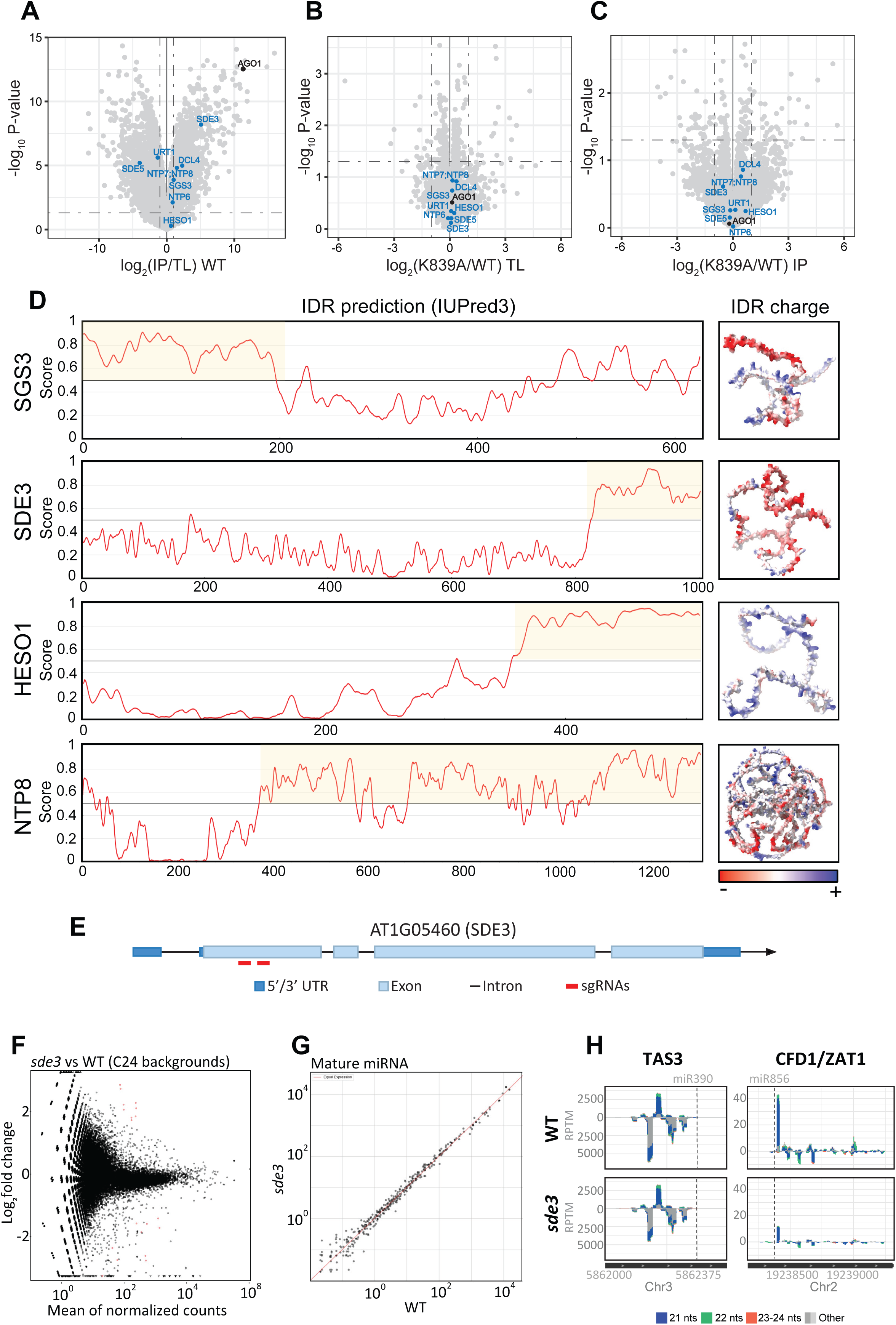
Characterization of candidate GW-interactors. **(A)** Volcano plot showing the differential abundance of proteins in total lysates (inflorescences) from FLAG-AGO1^K839A^ compared to FLAG-AGO1^WT^ plants. Protein abundance was determined by liquid chromatography-mass spectrometry. AGO1 is highlighted in black, and proteins relevant to this study are highlighted in light blue. **(B)** Volcano plot showing the differential abundance of proteins in total lysates *versus* co-purified fraction (inflorescences) from FLAG-AGO1^WT^ plants. Protein abundance was determined as in (A). AGO1 is highlighted in black, and proteins relevant to this study are highlighted in blue. **(C)** Volcano plot showing the differential abundance of proteins co-purified with FLAG-AGO1^K839A^ compared to FLAG-AGO1^WT^ from inflorescences. Protein abundance was determined by liquid chromatography-mass spectrometry of immunopurified fractions. AGO1 is highlighted in black, and proteins relevant to this study are highlighted in blue. **(D)** Protein disorder prediction for SGS3, SDE3, HESO1, and NTP8 using the IUPred3 web server. Intrinsically disordered regions (IDRs) are highlighted in yellow. Protein length is indicated below each plot. The right panels show structural views of the predicted IDRs generated by AlphaFold, with residues colored according to their charge on a red (negative) to blue (positive) scale. **(D)** SDE3 gene model with indication of the CRISPR-Cas9-generated deletion to knock out gene function. **(E)** MA plot showing differential accumulation of small RNAs from genes (red dots) in *sde3* and its corresponding wild-type (C24 background). **(F)** Scatter plots showing log₁₀-transformed reads per million (RPM) values of individual miRNAs detected in *sde3* plants compared to wild type (WT). The red diagonal line indicates equal abundance between the two backgrounds. **(G)** Small RNA accumulation in Reads Per Ten Million (RPTM) derived from *TAS3* and *CFD1* genes in inflorescences of *sde3* and C24 wild type (WT) plants. Gene model and genomic coordinates are shown below each plot (note: only partial gene models are displayed). miRNA target sites are highlighted with a dashed line. Small RNAs are color-coded by length: <21 nt (dark grey), 21 nt (blue), 22 nt (green), 23–24 nt (orange), and >24 nt (light grey).

**Figure EV6.**
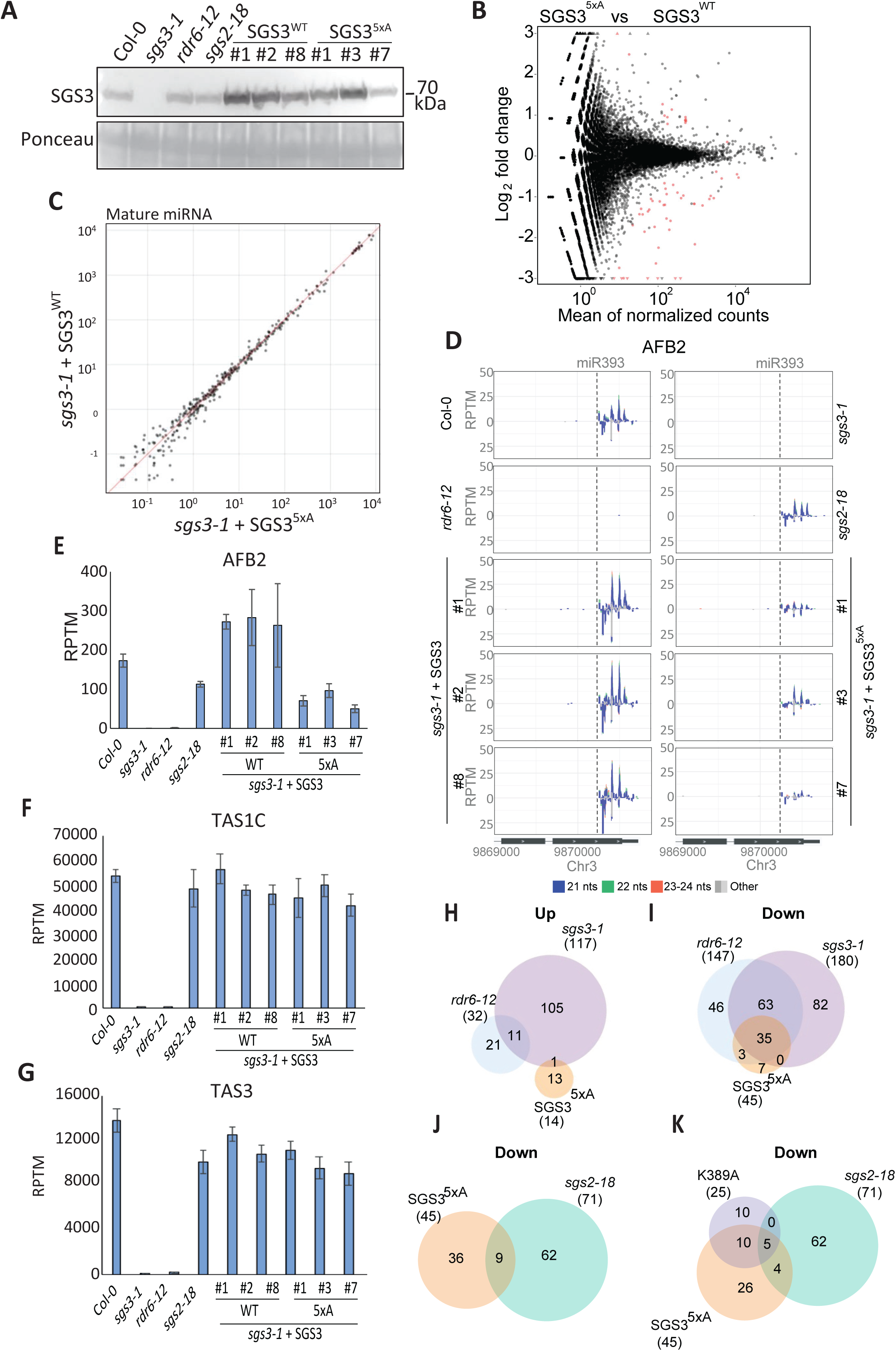
SGS3 IDR tryptophan residues are critical for siRNA amplification at multiple loci, but dispensable at others. **(A)** Immunoblot analysis of SGS3 levels in Col-0 WT, *sgs3-1*, *rdr6-12*, *sgs2-18*, and *sgs3-1* plants expressing SGS3^WT^ or SGS3^5xA^ (three independent biological replicates each), using total protein extracts from inflorescence tissue. Ponceau staining was used as a loading control. **(B)** MA plot showing differential accumulation of small RNAs from genes (red dots) in *sgs3-1* plants complemented with either SGS3^WT^ or SGS3^5xA^ versions. **(C)** Scatter plot showing log₁₀-transformed reads per million (RPM) values of individual miRNAs detected in *sgs3-1* plants complemented with either SGS3^WT^ or SGS3^5xA^ versions. The red diagonal line represents equal abundance between both genotypes. **(D)** Small RNA accumulation in Reads Per Ten Million (RPTM) derived from AFB2 gene in inflorescences of Col-0 WT*, sgs3-1, rdr6-12, sgs2-18 and sgs3-1* plants complemented with SGS3^WT^ or SGS3^5xA^ versions (three independent biological replicates of each). Gene model and genomic coordinates are shown below each plot (note: only partial gene models are displayed). miRNA target sites are highlighted with a dashed line. Small RNAs are color-coded by length: <21 nt (dark grey), 21 nt (blue), 22 nt (green), 23–24 nt (orange), and >24 nt (light grey). **(E)** Small RNA quantification of *AFB2*, *TAS1C* **(F)**, and *TAS3* **(G)** in the genotypes shown in (A). Bars represent the mean of three biological replicates, and error bars indicate the standard deviation (SD). Values are expressed as reads per ten million mapped reads (RPTM). **(H)-(I)** Venn diagrams illustrating the overlap of genes with up-(H) and downregulated (I) siRNAs between *sgs3-1*, *rdr6-12* and *sgs3-1* plants expressing SGS3^5xA^. Gene sets were derived from transcriptomic comparisons relative to Col-0 Wild-type (*sgs3-1* and *rdr6-12*), or *sgs3-1* plants complemented with SGS3^WT^ (SGS3^5xA^). **(J)** Venn diagram illustrating the overlap of genes with downregulated siRNAs between *sgs2-18* and *sgs3-1* plants expressing SGS3^5xA^. Gene sets were derived from transcriptomic comparisons relative to Col-0 WT for *sgs2-18*, and *sgs3-1* plants expressing SGS3^WT^ for SGS3^5xA^. **(K)** Venn diagram illustrating the overlap of genes with downregulated siRNAs between *sgs2-18*, *sgs3-1* expressing SGS3^5xA^, and *ago1-3* expressing FLAG-AGO1^K839^. Gene sets were derived from transcriptomic comparisons relative to Col-0 wild type for *sgs2-18), sgs3-1* plants expressing SGS3^WT^ for SGS3^5xA^, and *ago1-3* plants expressing FLAG-AGO1^WT^ for FLAG-AGO1^K839^.

**Figure EV7.**
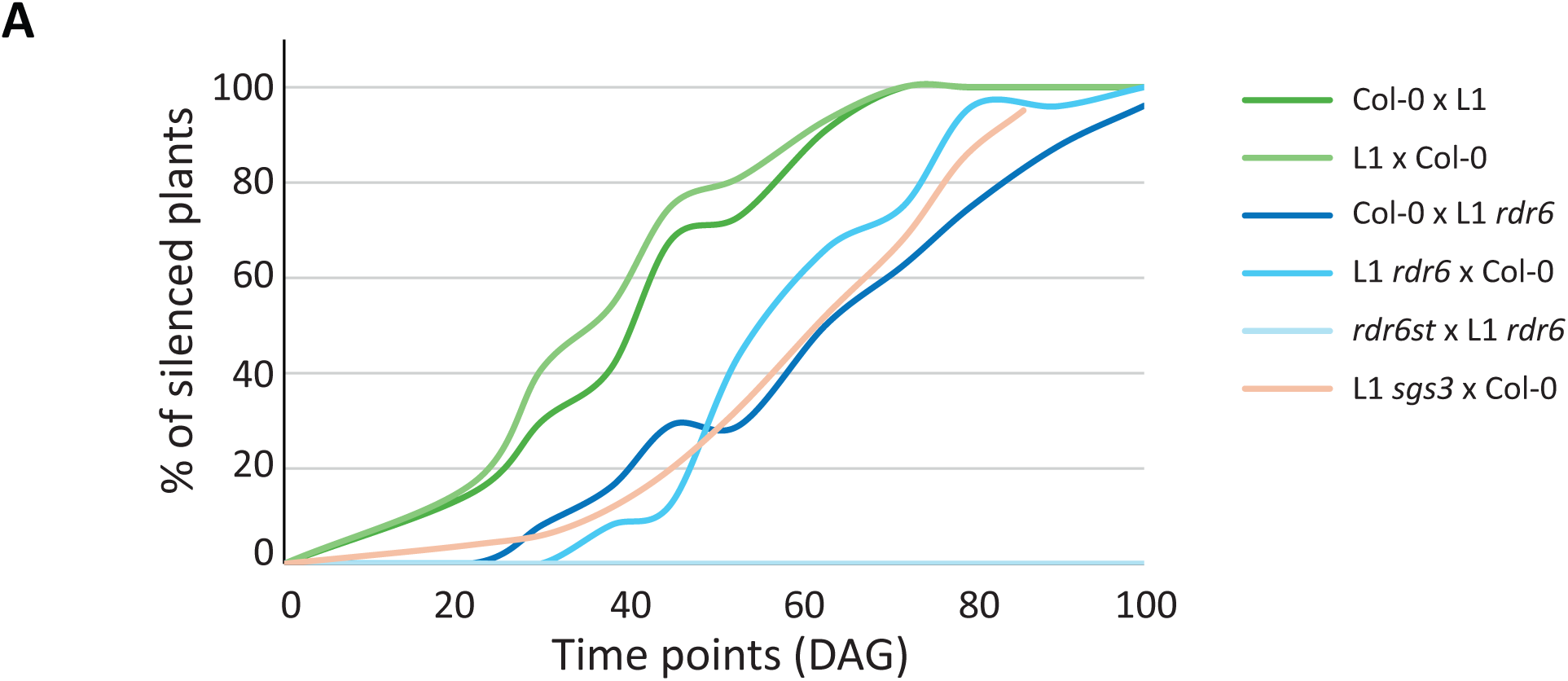
RDR6/SGS3 dosage determines the efficiency of *L1* transgene silencing. **(A)** Kinetics of the percentage of plants showing post-transcriptional gene silencing of GUS (*L1* line). Silencing was assessed by measuring GUS activity (arbitrary fluorescence units). Samples were collected from F1 populations of the indicated crosses at the specified days after germination.

**Figure EV8.**
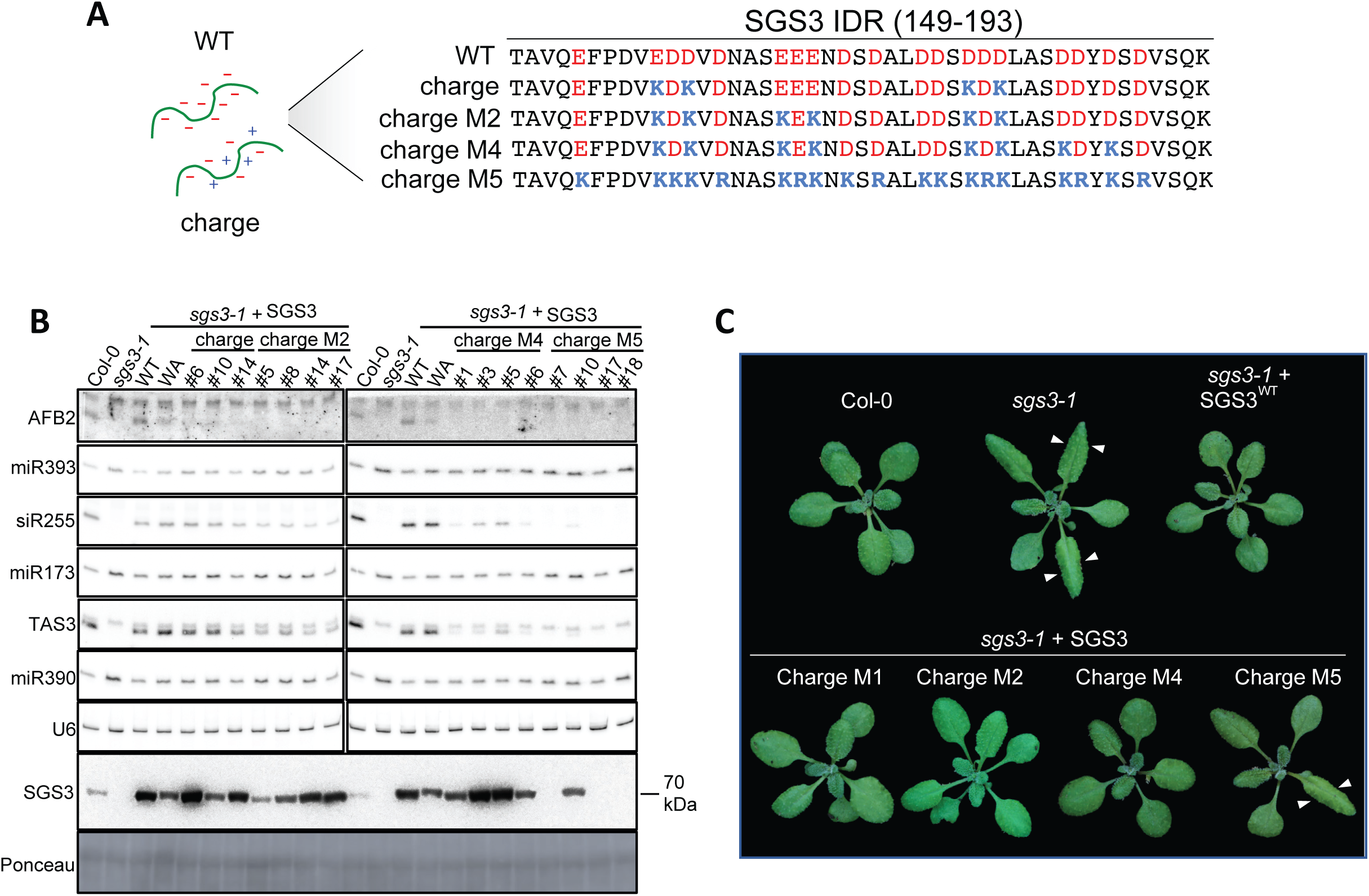
Progressive mutation of the SGS3 charged patch compromises tasiRNA biogenesis. **(A)** Partial amino acid sequence of the SGS3 IDR (amino acids 149-193) highlighting negatively charged residues (red) and those mutated to lysine or arginine (blue) in the different SGS3 charge mutant variants. **(B)** RNA blot analysis of secondary siRNAs and their corresponding trigger miRNAs in inflorescences of Col-0, *sgs3-1*, and *sgs3-1* plants expressing SGS3^WT^, SGS3^5xA^ or SGS3^charge^ variants. U6 serves as a loading control. SGS3 protein levels were analyzed by immunoblot from the same samples. Ponceau staining was used as a loading control. **(C)** Representative images of 3.5-week-old plants of Col-0 (WT), *sgs3-1*, and *sgs3-1* expressing SGS3^WT^ (Wild-type) or SGS3^charge^ variants (charge mutants). Arrows indicate the downward-curved, pointy leaves characteristic of *sgs3-1*.

## METHODS

### Plant material and growth conditions

All lines used in this study are in the *Arabidopsis thaliana* Col-0 background except the *sde3* mutant which is in the C24 background. The following mutants and transgenic lines have been described previously: *ago1-3* (Bohmert *et al*., 1998), *sgs3-1* (Mourrain *et al*., 2000), *rdr6-12* (Peragine *et al*., 2004), *ago1-3/FLAG-AGO1^WT^*(Arribas-Hernandez *et al*., 2016a), *35S:GUS* (*L1*) (Elmayan & Vaucheret, 1996b).

Arabidopsis seeds were surface-sterilized by washing for 2 minutes in 70% ethanol (v/v) followed by a wash step of 10 minutes in a solution containing 1.5% (w/v) sodium hypochlorite and 0.05% (w/v) Tween-20 and rinsed two times with sterile ddH2O. Then, seeds were plated in 1X Murashige and Skoog (MS) media supplemented with 1% sucrose and 0.8% agar and stratified in darkness for 72h at 4°C. Seedlings were grown under long day conditions (16h light/8h dark, 21°C, 100 mmol photons m^−2^s^−1^) in Aralab incubators. When needed, 12-day-old seedlings were transferred to soil (Plugg/Såjord [seed compost]; SW Horto) and grown under similar conditions (long-day photoperiod) in greenhouses or in a Percival/Aralab growth chamber.

### Transgenic line generation

*Arabidopsis thaliana* transgenic lines were generated by transformation of the corresponding backgrounds with *Agrobacterium tumefaciens* carrying the corresponding binary plasmid following the floral dipping method (Clough & Bent, 1998). Transformants were selected into MS plates supplemented with ampicillin (100 μg ml^-1^) and glufosinate-ammonium (BASTA, 7.5 μg ml^-1^), kanamycin (50 μg ml^-1^) or sulfadiazine (5 μg ml^-1^). The presence of the transgene was confirmed by PCR and/or detection of the corresponding proteins using western blot. At least two independent lines with a similar protein expression level and one single locus insertion were selected in T2, and homozygous T3 lines were identified when needed. After completed line selection, the *AGO1* transgenes were PCR amplified and sequenced to verify that the lines contained the correct AGO1 mutant variants.

### Cloning procedures

#### SGS3 variants

The full genomic fragment of SGS3, comprising ∼2 kb of the promoter region and 500 bp of the terminator region, was PCR-amplified from Arabidopsis genomic DNA using the KAPA HiFi Uracil+ Kit (Roche) and cloned into pLIFE41 by USER® cloning (New England Biolabs), following the manufacturer’s instructions. To generate SGS3 variants, DNA fragments containing the SGS3 IDR with SGS3^5xA^, SGS3^5xY^, or SGS3^charge^ modifications were synthesized by IDT, PCR-amplified with the KAPA HiFi Uracil+ Kit, and combined with the remaining SGS3 fragments and pLIFE41 (or derivatives) in a USER reaction. The sequences of all recombinant plasmids were verified by Sanger sequencing. Primers used for amplification are listed in Supplemental dataset 1.

#### Generation of FLAG-AGO1 variants by site directed mutagenesis

All FLAG-AGO1 variants were generated by inverted PCR using the pLIFE41-FLAG-AGO1^WT^ plasmid as a template (Arribas-Hernandez *et al*., 2016b). The sequences of all recombinant plasmids were verified by Sanger sequencing. Primers used for plasmid construction are listed in Supplemental dataset 1.

#### Plant genotyping

The *ago1-3* allele was identified as previously described (Arribas-Hernandez *et al*., 2016b). Briefly, primer LA298, which spans the region between the promoter and the coding sequence of the genomic *AGO1* locus and avoids amplification of any *AGO1* transgene, was used together with primer LA299 to amplify a 364-bp fragment from the genomic *AGO1* locus. After digestion with BseGI, the wild type fragment yields two bands of 324 and 40 bp, whereas three bands of 224, 100, and 40 bp are expected for the *ago1-3* allele.

The *sgs3-1* allele was identified by PCR amplification of a fragment from the endogenous *SGS3* locus using primers LA477 and LA478 (105 bp). After digestion with BsaI, the wild-type fragment yields a single band of 105 bp, whereas two bands of 85 and 20 bp are expected for the *sgs3-1* allele.

#### Generation of sde3 and mir173 deletions by CRISPR-Cas9

Knockout lines were generated using the pKIRT1.1 CRISPR-Cas9 system (Tsutsui & Higashiyama, 2017). Briefly, two plasmids were constructed for each target (*SDE3* or *MIR173*), each expressing an sgRNA generated by cloning two independent pairs of oligonucleotides (see Supplemental Dataset 1) targeting either the *SDE3* or *MIR173* locus. The plasmids were transformed into C24 for *sde3* or Col-0 for *mir173* plants, and transformants were selected on plates supplemented with hygromycin. Resistant seedlings were transferred to soil, and deletion mutants were identified by PCR using primers flanking the target regions (see Supplemental Dataset 1). Descendants of plants carrying deletions were counter-selected on hygromycin-containing plates to identify individuals that had lost the Cas9 transgene (hygromycin-sensitive). Plants lacking Cas9 were then transferred to fresh MS plates and recovered, and genotyped to identify individuals homozygous for the deletions of interest.

### Total RNA isolation

RNA extractions were performed using Tri Reagent (Thermofisher) following the manufacturer’s instructions. Biefly, 100 mg of flower tissue was flash-frozen and carefully ground using TissueLyser (Qiagen) or a mortar. 1 mL of Tri Reagent was added and well mixed with the flower powder by vortex. Cell debris was removed by centrifugation at 13,000 *g* for 10 minutes. The clarified plant lysate was transferred to a new Eppendorf tube and 0.2 ml of chloroform was added and mixed by vortex for 15 seconds. Then, phase separation was done by centrifugation at 4°C for 15 minutes at 13,000 *g* and the upper aqueous phase was transferred to a new tube. An equal volume of isopropanol was added, mixed by vortex and samples were incubated for 10 minutes at room temperature. RNA was precipitated by centrifugation at 13,000 *g* for 15 minutes at 4 °C and RNA pellets were washed two times with 75% (v/v) ethanol, dried and dissolved in RNAse-free water.

### Small RNA northern blot

Small RNA northern blots were performed as described (Arribas-Hernandez *et al*., 2016b). Briefly, 10-25 ug of total RNA was dissolved in 2x sRNA loading buffer (95% formamide, 18 mM EDTA pH 8.0, 0.025% sodium dodecyl sulfate (SDS), 0.01% bromophenol blue, 0.01% xylene cyanol) and denatured for 5 minutes at 95°C. RNAs were then resolved in a 0.5X TBE, 7M urea, 18% acrylamide gel run until bromophenol blue reached the bottom of the gel. RNA was then transferred to a Nylon membrane (Hybond-NX, Amersham) at 80V for 1 hour in cold 0.5X TBE. RNA was fixed to the membrane by chemical crosslinking in presence of EDC (1-ethyl-3-(3-dimethylaminopropyl) carbodiimide (Pall *et al*, 2007) for 1 hour at 60°C and membranes were washed two times with ddH_2_O and dried. After that, membranes were prehybridized for 30 minutes with PerfectHyb^TM^ Plus Hybridization buffer (Sigma) at 42°C and radiolabeled probes were added to the same buffer and incubated overnight at 42°C with gentle agitation. Membranes were then washed three times with 2x SSC, 2% SDS and exposed to a phosphorimager screen. Image acquisition was done using a Typhoon FLA 7000 scanner (GE Healthcare). When needed, membranes were stripped by treatment with 0.1% SDS boiling buffer three times.

For siR255, TAS3, miRNAs, and U6, complementary DNA oligonucleotides were 5′ end-labeled using T4 polynucleotide kinase (PNK; ThermoFisher) in the presence of γ^32^P-ATP. Probe sequences are provided in Supplemental dataset 1. For AFB2 and GUS, radioactively labeled probes were generated using PCR-amplified templates with the Prime-a-Gene labeling kit (Promega) in the presence of α^32^P-CTP, following the manufacturer’s instructions. Primer sequences used for amplification of probe templates are listed in Supplemental dataset 1.

### Protein blotting

Plant tissue was ground in the presence of 3x LDS and heated for 5 minutes at 90 °C and proteins were resolved by SDS-PAGE (Premade gels, BioRad). Then, proteins were blotted onto a Nitrocellulose membrane (Amershan Protan Premium) following the manufacturer’s instructions in a wet transfer system (BioRad). After that, membranes were staining with Ponceau and photographed as a loading control. Then, membranes were washed 5 minutes with 1x PBS with 0.05% tween-20 (v/v) and blocked 1 hour at room temperature in presence of 1 PBS, 5% non-fat milk (w/v) and 0.05% tween-20 (v/v). Afterwards, the primary antibody was added at the appropriate dilution and incubated at 4°C overnight with gentle rotation. The next day, membranes were washed three times, 10 minutes each, with 1X PBST, and incubated 1 hour at room temperature with the secondary antibody (when needed) in 1X PBS, 5% non-fat milk (w/v) and 0.05% Tween-20 (v/v). Membranes were washed three times at room temperature with 1X PBST and developed with a homemade ECL or using SuperSignal West Femto (Thermofisher).

### AGO1 immunoprecipitation

100 mg of ground flower tissue was resuspended into 0.4 mL of IP buffer (50 mM Tris HCL pH 7.5, 150 mM NaCl, 10% glycerol, 5 mM MgCl2, 0.1% NP40, 4 mM DTT and 1x EDTA free protease inhibitor) and incubated for15 minutes at 4°C with gentle agitation. The lysates were then centrifuged two times at 17,000 *g* for 10 minutes at 4°C to remove cell debris. After that, 10 µl of anti-flag agarose beads (20 ul slurry) were added and incubated for 1 hour at 4°C with rotation. Agarose beads were washed three times with IP wash buffer (IP buffer containing 500 mM NaCl). One half of the beads was dissolved in 1 mL of Tri Reagent to extract AGO1-bound RNAs following the manufacturer’s instruction, except that 20 µg of glycogen was added as a carrier at the RNA precipitation step. The AGO1-bound small RNA fraction was analyzed by northern blotting as described above. The remaining beads were dissolved in 40 ul of 1x LDS and heated for 5 minutes at 75°C to analyze AGO1 protein levels by western blot as explained below.

### Immunoprecipitation-mass spectrometry (IP-MS)

Immunoprecipitation of FLAG-AGO1 for mass spectrometry analysis was performed as described above, with minor modifications. Briefly, approximately 200 mg of ground flower tissue were resuspended in 0.8 mL of IP buffer (50 mM Tris-HCl pH 7.5, 150 mM NaCl, 10% glycerol, 5 mM MgCl₂, 0.1% NP-40, 4 mM DTT, and 1× EDTA-free protease inhibitor cocktail) and incubated for 5 minutes at 4°C with gentle agitation. The plant lysate was then centrifuged twice at 17,000 *g* for 10 minutes at 4°C to remove cell debris. Next, 10 µL of anti-FLAG agarose beads (Sigma) were added and incubated for 45 minutes at 4°C with rotation. Beads were washed twice with IP buffer (150 mM NaCl) and twice with IP buffer containing 2% glycerol. Protein elution was performed by incubating the beads with 60 µL of 100 ng/µL FLAG peptide (Sigma). Finally, 20% of the eluate was analyzed by SDS-PAGE followed by silver staining, and the remaining sample was analyzed by mass spectrometry following the procedure described below.

#### SP3 Desalting

Lysed samples were desalted using a standard SP3 (Single-Pot, Solid-Phase-Enhanced Sample Preparation) protocol. A 1:1 mixture of Sera-Mag™ Carboxylate-Modified Magnetic Beads & SpeedBeads (hydrophilic, GE45152105050250; hydrophobic, GE65152105050250; both 50 µg/µL stock) was prepared by vortexing each suspension vigorously and immediately extracting 120 µL of each to prevent bead settling. The two bead fractions were combined with 360 µL of MS-grade water, attached to a DynaMag-2™ magnetic rack, and the supernatant was removed after 1 min. Beads were resuspended in 1,000 µL MS-grade water, and this washing step was repeated three times before a final resuspension in 600 µL MS-grade water. For protein binding, 15 µL of washed beads were added to 300 µL of lysate, followed by 735 µL of 100% ethanol to achieve a final concentration of ∼70% (v/v). Samples were mixed gently by pipetting and incubated at 25°C for 15 min at 800 rpm using an Eppendorf ThermoMixer C. Bead–protein complexes were immobilized on the magnet for 1 min, the supernatant was removed, and beads were washed three times with 100 µL of 80% ethanol.

Proteins bound to the beads were digested overnight at 37 °C with shaking (1,000 rpm) in 50 µL of digestion buffer containing Trypsin/Lys-C protease mix (Pierce™ MS-grade, Thermo Scientific) at a protein:protease ratio of 10:1 (w/w).

#### EvoTip loading

Following digestion, samples were briefly pulse-spun to collect condensate prior to peptide clean-up using EvoTips (Evosep). The C18 material was activated with 20 µL of LiChrosolv® acetonitrile (ACN; Supelco, Cat. No. 1000292500) and centrifuged at 700 × g for 1 min. The tips were then submerged in LiChrosolv® 2-propanol (IPA; Supelco, Cat. No. 1027812500) for 1 min, followed by 20 µL of LiChrosolv® water containing 0.1% formic acid (Buffer A; Supelco, Cat. No. 1590132500), which was centrifuged through._Peptide digests were loaded into the activated tips and centrifuged, after which 20 µL of Buffer A was applied as a wash. Finally, 250 µL of Buffer A was added and centrifuged for 10 s at 700 × g. Approximately one-third of the EvoTip box was filled with Buffer A to prevent sample drying.

#### LC-MS analysis

Samples were analyzed using the Evosep One™ chromatography system (Evosep Biosystems) with a standard 40 SPD method, coupled to a Thermo Scientific™ Orbitrap™ Astral™ mass spectrometer equipped with a FAIMS Pro Duo interface (Thermo Scientific). Peptide separation was performed using an IonOpticks™ Aurora Elite™ TS C18 column (15 cm × 75 μm, 1.7 μm beads) maintained at 50 °C.

Data acquisition was performed using a WISH-DIA method (Petrosius *et al*, 2023), with MS1 spectra acquired at a resolution of 240,000 and a maximum injection time of 100 ms over a 400–800 m/z range, and MS2 spectra acquired over a 150–2000 m/z range with an HCD collision energy of 25%, a FAIMS compensation voltage of –48 V, an isolation window of 8 m/z, a total injection time of 30 ms, a loop control set to 35 scans, and an AGC target of 500% applied to both MS1 and MS2.

#### DIA search settings

TL and IP raw files were separately searched in DIA-NN (version 2.2.0) using the standard settings, with variable modifications for methionine oxidation and N-terminal acetylation enabled. The FASTA file used to generate the spectral library was the reviewed Arabidopsis thaliana UniProt protein database (18,719 entries), with NTP2 through NTP9 sequences added.

#### Data analysis and plot

Protein abundances were analyzed in R using the DEP (Zhang *et al*, 2018) workflow: log2 transformation, variance stabilizing normalization, and QRILC imputation of missing values. Principal component analysis (PCA) was used for sample inspection. Differential protein abundance was tested with limma linear models (paired for IP vs TL within WT; unpaired for K839A vs WT comparisons), with Benjamini–Hochberg FDR correction (Benjamini & Hochberg, 1995) and DEP significance cutoffs of log2 FC = 1 and adjusted p-value < 0.05. Results were visualized with volcano plots.

### In vitro binding assays

20 µl of streptavidin agarose beads (Solulink) were incubated with 20 µg of biotinylated peptides in 40 µl of PBS for 30 min at RT under constant agitation. Beads were washed three times with 1 ml PBS to remove excess peptide and incubated for 2h at 4°C with 500 µl of plant lysate prepared from 250 mg of inflorescence tissue. The beads were then washed twice with 1 ml PBS, and the supernatant was removed. Proteins retained on the beads were extracted with 30 µl of Laemmli buffer, heated to 85 °C for 5 min, and subjected to SDS–PAGE. Western blotting was performed using an anti-FLAG antibody (Sigma) at 1:750 dilution and developed with SuperSignal™ West Femto Maximum Sensitivity Substrate (Life Technologies).

### GUS extraction and activity quantification

GUS extraction and activity quantification were performed as previously described (Elmayan & Vaucheret, 1996a). Briefly, soluble protein fractions were extracted from cauline leaves of flowering plants using phosphate buffer (50 mM NaPO₄ pH 7.0, 10 mM Na₂EDTA, and 40 mM β-mercaptoethanol). Equal amounts of total protein (1–4 µg) were incubated with 1 mM 4-methylumbelliferyl-β-D-glucuronide (Duchefa), and GUS activity was quantified by measuring the amount of 4-methylumbelliferone produced using a fluorometer (Fluoroscan II; Thermo Scientific).

### Small RNA library preparation and sequencing

AGO-bound small RNAs were purified using the TrapR method (Grentzinger *et al*., 2020). TrapR columns were prepared using Q Sepharose HP resin (GE healthcare, GE17-1014-01) as described (Grentzinger *et al*., 2020). In brief, 30 mg of flower tissue was resuspended in 400 uL of lysis buffer (20 mM HEPES–KOH, pH 7.9, 10% (v/v) glycerol, 1.5 mM MgCl2, 0.2 mM EDTA, 1 mM DTT, 0.1% (v/v) Triton X-100, conductivity adjusted at 8 mS cm^−2^ with KOAc. Then, the plant lysate was centrifuged 5 minutes at 10000g at 4°C to remove cell debris. The cleared lysate was then transferred to a new Eppendorf tube and 1/4 of the sample (100 ul) was saved for analysis of the input. Columns were centrifuged 30 seconds at 1000 *g* to remove the storage buffer and 300 uL of the samples were added to the columns. The lysate and the resin were mixed, and columns centrifuged 30 seconds at 1000 *g* to collect the elution fraction. Then, 300 ul of the TrapR elution buffer (20 mM HEPES– KOH, pH 7.9, 10% (v/v) glycerol, 1.5 mM MgCl2, 0.2 mM EDTA, 1 mM DTT, conductivity adjusted at 40 mS cm^−2^ with KOAc) was added, mixed and centrifuged for 30 seconds at 1000 *g* to collect again the “Elution-fraction”. This step was repeated once (900 ul of “Elution fraction”). The TrapR purified RNAs were extracted by mixing with 1 vol (900 ul) of Roti-Aqua-P/C/I pH 4.5–5 (Carl Roth). The aqueous phase was isolated by centrifugation at 14,000 rpm for 15 minutes at 4°C and transferred to a new Eppendorf tube. RNA precipitation was performed in presence of 1/10 vol of 3M sodium acetate (pH 5.3), 1 µl of glycogen (Thermofisher) and 1 vol of isopropanol by overnight incubation at -20°C followed by centrifugation at 14,000 rpm for 30 minutes. RNA was resuspended in 12 µl of RNAse-free water.

Before small RNA library preparation, 1 ul of the purified small RNA fraction was end-labelled with PNK in an exchange reaction (1x PNK Buffer B, Thermofisher) in the presence of ^32^P-γ-ATP, and resolved on a 7M urea, 18% polyacrylamide (19:1) gel to check small RNA integrity. Small RNA libraries were prepared using the NEBNext Multiplex Small RNA Library Prep Set (#E7300S, New England Biolabs) following the manufacturer’s instructions.

cDNA libraries were analyzed using an Agilent Bioanalyzer, pooled together and size-selected on a 6% polyacrylamide gel. The purified size-selected pool of libraries was then quantified using Qubit (Invitrogen) and single-end sequenced on an Illumina Nextseq 500 with SE75 HI (Illumina).

### Processing of sRNA sequencing and bioinformatic analysis

Raw reads were first quality-inspected using FASTQC (Andrews, 2010). Trimming of the reads was done using Trimmomatic (Bolger *et al*, 2014). Afterwards, quality was confirmed using FASTQC, and the reads mapped to Arabidopsis genome (TAIR10 version) using Bowtie (Langmead *et al*, 2009) with no mismatch allowed (-v 0 mode) except for the *sde3* sequencing (C24 background, -v 1 mode). SAM files were converted to BAM files, sorted and indexed using samtools (Li *et al*, 2009). Mapping of the reads and plot generation was visualized using the IGV browser (Robinson *et al*, 2011) or sRNA Viewer tool (Michael J. Axtell; https://github.com/MikeAxtell/sRNA_Viewer). The number of reads per feature were estimated using HTSeq-count (Anders *et al*, 2014). Statistical analyses were performed using DESeq2 (Love *et al*, 2014a). Mature miRNA profiles were generated by retrieving all mapped reads corresponding to mature miRNAs (TAIR10 GFF annotation). Reads per million (RPM) were calculated using the total number of mapped reads as a normalization factor. Plots were generated in R using in-house scripts. For Venn diagrams, overlapping gene sets were calculated with InteractiVenn (Heberle *et al*, 2015), and final plots were produced with the venneuler R package. Total miRNA counts, small RNA size distributions, and 5′-nucleotide compositions were estimated using in-house python/R scripts. MA plots, hierarchical clustering dendrograms, and principal component analysis (PCA) plots were generated with DESeq2 (Love *et al*, 2014b).

### IDR predictions and visualization

Intrinsic disorder predictions for full-length SGS3, SDE3, HESO1, and NTP8 proteins were performed using IUPred3 (Erdős *et al*, 2021). Predicted IDRs were subsequently visualized with UCSF ChimeraX (Meng *et al*, 2023), where residues were colored according to their charge.

### SGS3 IDR analysis

Protein sequences in FASTA format were screened for conserved motifs at the beginning of the globular domain of SGS3 (“KWFK” or “RWFK”). Sequences without either motif were no further analyzed. For each retained sequence, the intrinsically disordered region (IDR) was defined as the segment from the first residue up to (but not including) the first occurrence of the motif; these IDRs were written to a new FASTA file (N = 268). For the metaplot, each IDR of length L was mapped to the unit interval [0,1] by dividing it into N = 100 equal-width bins using rounded integer boundaries (bin k spans indices round(kL/N) to round((k+1)L/N)−1). Within each bin, the per-sequence density of amino acid *A* was computed as the fraction of residues of type *A* in that bin: density = (#*A* in bin / bin size). Densities were averaged across IDRs (equal weight per sequence) to obtain mean profiles along the normalized IDR length for R, K, E, D and W. The resulting profiles were Gaussian-smoothed (σ = 3 bins) and plotted with Matplotlib. Using the same IDR set, we also quantified the number of tryptophans (W) per IDR, the sequence context surrounding tryptophans, and the distribution of IDR lengths.

### Structural analysis and conservation of Argonaute proteins

Structural visualization and comparative analysis were carried out using PyMOL (v3.1.6.1; Schrödinger, LLC). The x-ray structure of human Argonaute 2 (HsAgo2; PDB ID: 6CBD) was used as a reference for the structural analysis. *Arabidopsis thaliana* Argonaute proteins (AtAgo1-10) and AtAgo1 R834E and R1045E were modelled using the AlphaFold v3 prediction (https://alphafoldserver.com/) in an RNA free state. The tryptophan binding pockets 1 and 3 from the HsAgo2 structure served as reference points for the structural alignment and interpretation of their structural conservation to plants.

The conservation of surface charge was examined by individual calculations of the electrostatic potential using the Adaptive Poisson-Boltzmann Solver (APBS) plugin in PyMOL using default parameters and range setting of +/-3.0. All molecular representations, electrostatic surface renderings were generated in PyMOL for figure preparation. Standard primary sequence alignments were generated by Clustal Omega (Sievers *et al*, 2011) and a phylogenetic tree of AtAgo1-10 was built by the Neighbor-Joining method.

## DATA AVAILABILITY

High-throughput sequencing data have been deposited in the Sequence Read Archive (SRA; NCBI) under the BioProject ID PRJNA1327840. Mass spectrometry data have been deposited at the Proteomics Identification Database (PRIDE) (https://www.ebi.ac.uk/pride/) under project accession PXD068382.

## Reviewer Access To Mass Spectrometry Data In Pride

Log in to the PRIDE website using the following details:

**Project accession:** PXD068382

**Token:** 48t2KdAwmhps

Alternatively, access the dataset by logging in to the PRIDE website using the following account details:

**Password:** 4qElQXHL6uK2

## CONFLICT OF INTEREST

The authors declare that they have no conflict of interest.

## ACKNOWLEDGMENTS

Freja Asmussen is thanked for constructing CRISPR mutants in the *MIR173* gene and Juliane Birklund Wezelenburg for assistance with the *sde3* CRISPR mutant construction. Theo Bölsterli and Rene Hvidberg Petersen and their teams are thanked for plant care, and Lena Bjørn Johansson for valuable technical assistance. Sébastien Bélanger and Blake Meyers are thanked for providing sequences of plant SBS3 orthologs. This work was supported by research grants from Novo Nordisk Foundation (Hallas Møller 2010), European Research Council (ERC-2011-StG 282460 and ERC-2016-CoG 726417), Lundbeck Foundation (R83-A7859), and Villum Fonden (13397), and by infrastructure grants from Carlsberg Fondet (CF18-1075 and CF20-0659), Augustinus Fonden (12-5126) and Brdr Hartmann Fonden (A35879), all to P.B.

## AUTHOR CONTRIBUTIONS

**Diego López-Márquez:** Conceptualization; Investigation, Formal analysis, Supervision; Visualization; Writing-original draft; Writing-review and editing

**Laura Arribas-Hernández:** Conceptualization; Investigation, Writing-review and editing

**Christian Poulsen:** Conceptualization; Visualization, Writing-review and editing

**Emilie Duus Oksbjerg:** Investigation; Writing-review and editing

**Nathalie Bouteiller:** Investigation

**Mads Meier:** Investigation

**Johannes Blanke:** Investigation; Writing-review and editing

**Angel del Espino:** Investigation

**Maria Louisa Vigh:** Investigation, Writing-review and editing

**Simon Bressendorff:** Formal analysis; Investigation; Writing-review and editing

**Alberto Carbonell:** Investigation; Writing-review and editing

**Rune Daucke:** Investigation; Formal analysis; Writing-review and editing

**Erwin M. Schoof:** Supervision; Writing-review and editing

**Hervé Vaucheret:** Investigation; Supervision; Formal analysis; Conceptualization; Writing-review and editing

**Peter Brodersen:** Conceptualization; Supervision; Funding acquisition; Data curation; Visualization; Methodology; Project administration, Writing-original draft, Writing-review and editing

## Notes

### Competing Interest Statement

The authors have declared no competing interest.

## REFERENCES

1. Adenot X, Elmayan T, Lauressergues D, Boutet S, Bouche N, Gasciolli V, Vaucheret H (2006) DRB4-dependent TAS3 trans-acting siRNAs control leaf morphology through AGO7. Current biology : CB 16: 927–932

2. Allen E, Xie Z, Gustafson AM, Carrington JC (2005) microRNA-directed phasing during trans-acting siRNA biogenesis in plants. Cell 121: 207–221

3. Anders S, Pyl PT, Huber W (2014) HTSeq—a Python framework to work with high-throughput sequencing data. Bioinformatics 31: 166–169

4. Andrews S (2010) FastQC: A Quality Control Tool for High Throughput Sequence Data *Available online*; http://www.bioinformaticsbabrahamacuk/projects/fastqc/

5. Arribas-Hernandez L, Kielpinski LJ, Brodersen P (2016a) mRNA Decay of Most Arabidopsis miRNA Targets Requires Slicer Activity of AGO1. Plant physiology 171: 2620–2632

6. Arribas-Hernandez L, Marchais A, Poulsen C, Haase B, Hauptmann J, Benes V, Meister G, Brodersen P (2016b) The Slicer Activity of ARGONAUTE1 Is Required Specifically for the Phasing, Not Production, of Trans-Acting Short Interfering RNAs in Arabidopsis. The Plant cell 28: 1563–1580

7. Axtell MJ, Jan C, Rajagopalan R, Bartel DP (2006) A two-hit trigger for siRNA biogenesis in plants. Cell 127: 565–577

8. Baumberger N, Baulcombe DC (2005) Arabidopsis ARGONAUTE1 is an RNA Slicer that selectively recruits microRNAs and short interfering RNAs. Proceedings of the National Academy of Sciences of the United States of America 102: 11928–11933

9. Bélanger S, Zhan J, Meyers BC (2023) Phylogenetic analyses of seven protein families refine the evolution of small RNA pathways in green plants. Plant physiology 192: 1183–1203

10. Benjamini Y, Hochberg Y (1995) Controlling the False Discovery Rate: A Practical and Powerful Approach to Multiple Testing. Journal of the Royal Statistical Society: Series B (Methodological) 57: 289–300

11. Bies-Etheve N, Pontier D, Lahmy S, Picart C, Vega D, Cooke R, Lagrange T (2009) RNA-directed DNA methylation requires an AGO4-interacting member of the SPT5 elongation factor family. EMBO reports 10: 649–654

12. Bohmert K, Camus I, Bellini C, Bouchez D, Caboche M, Benning C (1998) AGO1 defines a novel locus of Arabidopsis controlling leaf development. EMBO Journal 17: 170–180

13. Bolger AM, Lohse M, Usadel B (2014) Trimmomatic: a flexible trimmer for Illumina sequence data. Bioinformatics 30: 2114–2120

14. Branscheid A, Marchais A, Schott G, Lange H, Gagliardi D, Andersen SU, Voinnet O, Brodersen P (2015) SKI2 mediates degradation of RISC 5’-cleavage fragments and prevents secondary siRNA production from miRNA targets in Arabidopsis. Nucleic acids research 43: 10975–10988

15. Braun JE, Huntzinger E, Izaurralde E (2013) The Role of GW182 Proteins in miRNA-Mediated Gene Silencing. In: Ten Years of Progress in GW/P Body Research, Chan E.K.L., Fritzler M.J. (eds.) pp. 147–163. Springer New York: New York, NY

16. Bressendorff S, Sjøgaard IMZ, Prestel A, Voutsinos V, Jansson MD, Ménard P, Lund AH, Hartmann-Petersen R, Kragelund BB, Poulsen C et al (2025) Importance of an N-terminal structural switch in the distinction between small RNA-bound and free ARGONAUTE. Nature structural & molecular biology 32: 625–638

17. Briskin D, Wang PY, Bartel DP (2020) The biochemical basis for the cooperative action of microRNAs. Proceedings of the National Academy of Sciences 117: 17764–17774

18. Brodersen P, Sakvarelidze-Achard L, Bruun-Rasmussen M, Dunoyer P, Yamamoto YY, Sieburth L, Voinnet O (2008) Widespread translational inhibition by plant miRNAs and siRNAs. Science 320: 1185–1190

19. Bukhari SIA, Truesdell SS, Lee S, Kollu S, Classon A, Boukhali M, Jain E, Mortensen RD, Yanagiya A, Sadreyev RI et al (2016) A Specialized Mechanism of Translation Mediated by FXR1a-Associated MicroRNP in Cellular Quiescence. Molecular cell 61: 760–773

20. Carbonell A, Fahlgren N, Garcia-Ruiz H, Gilbert KB, Montgomery TA, Nguyen T, Cuperus JT, Carrington JC (2012) Functional analysis of three Arabidopsis ARGONAUTES using slicer-defective mutants. The Plant cell 24: 3613–3629

21. Chen H-M, Chen L-T, Patel K, Li Y-H, Baulcombe DC, Wu S-H (2010) 22-nucleotide RNAs trigger secondary siRNA biogenesis in plants. Proceedings of the National Academy of Sciences 107: 15269–15274

22. Cisneros Adriana E, Alarcia A, Llorens-Gámez Juan J, Puertes A, Juárez-Molina M, Primc A, Carbonell A (2025) Syn-tasiR-VIGS: virus-based targeted RNAi in plants by synthetic trans-acting small interfering RNAs derived from minimal precursors. Nucleic acids research 53

23. Clough SJ, Bent AF (1998) Floral dip: a simplified method for Agrobacterium-mediated transformation of Arabidopsis thaliana. Plant J 16: 735–743

24. Cuperus JT, Carbonell A, Fahlgren N, Garcia-Ruiz H, Burke RT, Takeda A, Sullivan CM, Gilbert SD, Montgomery TA, Carrington JC (2010) Unique functionality of 22-nt miRNAs in triggering RDR6-dependent siRNA biogenesis from target transcripts in Arabidopsis. Nature structural & molecular biology 17: 997–1003

25. Dalmay T, Hamilton AJ, Rudd S, Angell S, Baulcombe DC (2000) An RNA-dependent RNA polymerase gene in *Arabidopsis* is required for posttranscriptional gene silencing mediated by a transgene but not by a virus. Cell 101: 543–553

26. Dalmay TD, Horsefield R, Braunstein TH, Baulcombe DC (2001) SDE3 encodes an RNA helicase required for post-transcriptional gene silencing in Arabidopsis. EMBO Journal 20: 2069–2078

27. de Felippes Felipe F, Marchais A, Sarazin A, Oberlin S, Voinnet O (2017) A single miR390 targeting event is sufficient for triggering TAS3-tasiRNA biogenesis in Arabidopsis. Nucleic acids research 45: 5539–5554

28. Derrien B, Clavel M, Baumberger N, Iki T, Sarazin A, Hacquard T, Ponce MR, Ziegler-Graff V, Vaucheret H, Micol JL et al (2018) A Suppressor Screen for AGO1 Degradation by the Viral F-Box P0 Protein Uncovers a Role for AGO DUF1785 in sRNA Duplex Unwinding. The Plant cell 30: 1353–1374

29. Doench JG, Sharp PA (2004) Specificity of microRNA target selection in translational repression. Genes & development 18: 504–511

30. Dunoyer P, Himber C, Voinnet O (2005) DICER-LIKE 4 is required for RNA interference and produces the 21-nucleotide small interfering RNA component of the plant cell-to-cell silencing signal. Nature genetics 37: 1356–1360

31. El-Shami M, Pontier D, Lahmy S, Braun L, Picart C, Vega D, Hakimi MA, Jacobsen SE, Cooke R, Lagrange T (2007) Reiterated WG/GW motifs form functionally and evolutionarily conserved ARGONAUTE-binding platforms in RNAi-related components. Genes & development 21: 2539–2544

32. Elkayam E, Faehnle CR, Morales M, Sun J, Li H, Joshua-Tor L (2017) Multivalent Recruitment of Human Argonaute by GW182. Molecular cell 67: 646–658.e643

33. Elmayan T, Adenot X, Gissot L, Lauressergues D, Gy I, Vaucheret H (2009) A neomorphic sgs3 allele stabilizing miRNA cleavage products reveals that SGS3 acts as a homodimer. The FEBS Journal 276: 835–844

34. Elmayan T, Vaucheret H (1996a) Expression of single copies of a strongly expressed 35S transgene can be silenced post-transcriptionally. The Plant Journal 9: 787–797

35. Elmayan T, Vaucheret H (1996b) Expression of single copies of a strongly expressed *35S* transgene can be silenced post-transcriptionally. Plant Journal 9: 787–797

36. Erdős G, Pajkos M, Dosztányi Z (2021) IUPred3: prediction of protein disorder enhanced with unambiguous experimental annotation and visualization of evolutionary conservation. Nucleic acids research 49: W297–W303

37. Eulalio A, Helms S, Fritzsch C, Fauser M, Izaurralde E (2009) A C-terminal silencing domain in GW182 is essential for miRNA function. Rna 15: 1067–1077

38. Eulalio A, Huntzinger E, Izaurralde E (2008) GW182 interaction with Argonaute is essential for miRNA-mediated translational repression and mRNA decay. Nature structural & molecular biology 15: 346–353

39. Fagard M, Boutet S, Morel J-B, Bellini C, Vaucheret H (2000) AGO1, QDE-2, and RDE-1 are related proteins required for post-transcriptional gene silencing in plants, quelling in fungi, and RNA interference in animals. Proc Natl Acad Sci USA 97: 11650–11654

40. Fahlgren N, Montgomery TA, Howell MD, Allen E, Dvorak SK, Alexander AL, Carrington JC (2006) Regulation of AUXIN RESPONSE FACTOR3 by TAS3 ta-siRNA Affects Developmental Timing and Patterning in Arabidopsis. Curr Biol 16: 939–944

41. Fang X, Qi Y (2016) RNAi in Plants: An Argonaute-Centered View. The Plant cell 28: 272–285

42. Fei Q, Yu Y, Liu L, Zhang Y, Baldrich P, Dai Q, Chen X, Meyers BC (2018) Biogenesis of a 22-nt microRNA in Phaseoleae species by precursor-programmed uridylation. Proceedings of the National Academy of Sciences 115: 8037–8042

43. Felippes FF, Weigel D (2009) Triggering the formation of tasiRNAs in Arabidopsis thaliana: the role of microRNA miR173. EMBO reports 10: 264–270

44. Fukunaga R, Doudna JA (2009) dsRNA with 5′ overhangs contributes to endogenous and antiviral RNA silencing pathways in plants. The EMBO journal 28: 545–555

45. Garcia D, Collier SA, Byrne ME, Martienssen RA (2006) Specification of leaf polarity in Arabidopsis via the trans-acting siRNA pathway. Current biology : CB 16: 933–938

46. Garcia D, Garcia S, Pontier D, Marchais A, Renou JP, Lagrange T, Voinnet O (2012) Ago hook and RNA helicase motifs underpin dual roles for SDE3 in antiviral defense and silencing of nonconserved intergenic regions. Molecular cell 48: 109–120

47. Garcia-Ruiz H, Takeda A, Chapman EJ, Sullivan CM, Fahlgren N, Brempelis KJ, Carrington JC (2010) Arabidopsis RNA-dependent RNA polymerases and Dicer-like proteins in antiviral defense and small interfering RNA biogenesis during Turnip Mosaic Virus infection. The Plant cell 22: 481–496

48. Gasciolli V, Mallory AC, Bartel DP, Vaucheret H (2005) Partially Redundant Functions of Arabidopsis DICER-like Enzymes and a Role for DCL4 in Producing trans-Acting siRNAs. Current biology : CB 16: 1494–1500

49. Grentzinger T, Oberlin S, Schott G, Handler D, Svozil J, Barragan-Borrero V, Humbert A, Duharcourt S, Brennecke J, Voinnet O (2020) A universal method for the rapid isolation of all known classes of functional silencing small RNAs. Nucleic acids research 48: e79–e79

50. Heberle H, Meirelles GV, da Silva FR, Telles GP, Minghim R (2015) InteractiVenn: a web-based tool for the analysis of sets through Venn diagrams. BMC Bioinformatics 16: 169

51. Himber C, Dunoyer P, Moissiard G, Ritzenthaler C, Voinnet O (2003) Transitivity-dependent and -independent cell-to-cell movement of RNA silencing. The EMBO journal 22: 4523–4533

52. Howell MD, Fahlgren N, Chapman EJ, Cumbie JS, Sullivan CM, Givan SA, Kasschau KD, Carrington JC (2007) Genome-wide analysis of the RNA-DEPENDENT RNA POLYMERASE6/DICER-LIKE4 pathway in Arabidopsis reveals dependency on miRNA-and tasiRNA-directed targeting. The Plant cell 19: 926–942

53. Hunter C, Willmann MR, Wu G, Yoshikawa M, de la Luz Gutierrez-Nava M, Poethig SR (2006) Trans-acting siRNA-mediated repression of ETTIN and ARF4 regulates heteroblasty in Arabidopsis. Development 133: 2973–2981

54. Iwakawa H-o, Lam AYW, Mine A, Fujita T, Kiyokawa K, Yoshikawa M, Takeda A, Iwasaki S, Tomari Y (2021) Ribosome stalling caused by the Argonaute-microRNA-SGS3 complex regulates the production of secondary siRNAs in plants. Cell reports 35

55. Jannot G, Michaud P, Quévillon Huberdeau M, Morel-Berryman L, Brackbill JA, Piquet S, McJunkin K, Nakanishi K, Simard MJ (2016) GW182-Free microRNA Silencing Complex Controls Post-transcriptional Gene Expression during Caenorhabditis elegans Embryogenesis. PLoS genetics 12: e1006484

56. Jauvion V, Elmayan T, Vaucheret H (2010) The Conserved RNA Trafficking Proteins HPR1 and TEX1 Are Involved in the Production of Endogenous and Exogenous Small Interfering RNA in Arabidopsis The Plant cell 22: 2697–2709

57. Jouannet V, Moreno AB, Elmayan T, Vaucheret H, Crespi MD, Maizel A (2012) Cytoplasmic Arabidopsis AGO7 accumulates in membrane-associated siRNA bodies and is required for ta-siRNA biogenesis. The EMBO journal 31: 1704–1713

58. Kim G-T, Tsukaya H, Uchimiya H (1998) The *ROTUNDIFOLIA3* gene of *Arabidopsis thaliana* encodes a new member of the cytochrome P-450 family that is required for the regulated polar elongation of leaf cells. Genes & development 12: 2381–2391

59. Kuzuoğlu-Öztürk D, Bhandari D, Huntzinger E, Fauser M, Helms S, Izaurralde E (2016) miRISC and the CCR4-NOT complex silence mRNA targets independently of 43S ribosomal scanning. The EMBO journal 35: 1186–1203

60. Lahmy S, Pontier D, Bies-Etheve N, Laudié M, Feng S, Jobet E, Hale CJ, Cooke R, Hakimi M-A, Angelov D et al (2016) Evidence for ARGONAUTE4–DNA interactions in RNA-directed DNA methylation in plants. Genes & development 30: 2565–2570

61. Langmead B, Trapnell C, Pop M, Salzberg SL (2009) Ultrafast and memory-efficient alignment of short DNA sequences to the human genome. Genome biology 10: R25

62. Li H, Handsaker B, Wysoker A, Fennell T, Ruan J, Homer N, Marth G, Abecasis G, Durbin R, Genome Project Data Processing S (2009) The Sequence Alignment/Map format and SAMtools. Bioinformatics 25: 2078–2079

63. Li S, Liu L, Zhuang X, Yu Y, Liu X, Cui X, Ji L, Pan Z, Cao X, Mo B et al (2013) MicroRNAs inhibit the translation of target mRNAs on the endoplasmic reticulum in Arabidopsis. Cell 153: 562–574

64. Liu J, Carmell MA, Rivas FV, Marsden CG, Thomson JM, Song JJ, Hammond SM, Joshua-Tor L, Hannon GJ (2004) Argonaute2 is the catalytic engine of mammalian RNAi. Science 305: 1437–1441

65. Liu Y, Teng C, Xia R, Meyers BC (2020) PhasiRNAs in Plants: Their Biogenesis, Genic Sources, and Roles in Stress Responses, Development, and Reproduction. The Plant cell 32: 3059–3080

66. Love MI, Huber W, Anders S (2014a) Moderated estimation of fold change and dispersion for RNA-seq data with DESeq2. Genome biology 15: 550

67. Love MI, Huber W, Anders S (2014b) Moderated estimation of fold change and dispersion for RNA-seq data with DESeq2. Genome biology 15: 550

68. Ma JB, Ye K, Patel DJ (2004) Structural basis for overhang-specific small interfering RNA recognition by the PAZ domain. Nature 429: 318–322

69. Ma JB, Yuan YR, Meister G, Pei Y, Tuschl T, Patel DJ (2005) Structural basis for 5’-end-specific recognition of guide RNA by the A. fulgidus Piwi protein. Nature 434: 666–670

70. Manavella PA, Koenig D, Weigel D (2012) Plant secondary siRNA production determined by microRNA-duplex structure. Proceedings of the National Academy of Sciences of the United States of America 109: 2461–2466

71. Martin EW, Holehouse AS, Peran I, Farag M, Incicco JJ, Bremer A, Grace CR, Soranno A, Pappu RV, Mittag T (2020) Valence and patterning of aromatic residues determine the phase behavior of prion-like domains. Science 367: 694–699

72. Matsuura-Suzuki E, Kiyokawa K, Iwasaki S, Tomari Y (2024) miRNA-mediated gene silencing in *Drosophila* larval development involves GW182-dependent and independent mechanisms. The EMBO journal 43: 6161–6179

73. Meister G, Landthaler M, Patkaniowska A, Dorsett Y, Teng G, Tuschl T (2004) Human Argonaute2 mediates RNA cleavage targeted by miRNAs and siRNAs. Molecular cell 15: 185–197

74. Meng EC, Goddard TD, Pettersen EF, Couch GS, Pearson ZJ, Morris JH, Ferrin TE (2023) UCSF ChimeraX: Tools for structure building and analysis. Protein Science 32: e4792

75. Mi S, Cai T, Hu Y, Chen Y, Hodges E, Ni F, Wu L, Li S, Zhou H, Long C et al (2008) Sorting of small RNAs into Arabidopsis argonaute complexes is directed by the 5’ terminal nucleotide. Cell 133: 116–127

76. Mittag T, Pappu RV (2022) A conceptual framework for understanding phase separation and addressing open questions and challenges. Molecular cell 82: 2201–2214

77. Morel J-B, Gordon C, Mourrain P, Beclin C, Boutet S, Feuerbach F, Proux F, Vaucheret H (2002) Fertile hypomorphic *ARGONAUTE* (*ago1*) mutants impaired in post-transcriptional gene silencing and virus resistance. The Plant cell 14: 629–639

78. Motamedi MR, Verdel A, Colmenares SU, Gerber SA, Gygi SP, Moazed D (2004) Two RNAi complexes, RITS and RDRC, physically interact and localize to noncoding centromeric RNAs. Cell 119: 789–802

79. Mourrain P, Beclin C, Elmayan T, Feuerbach F, Godon C, Morel J-B, Jouette D, Lacombe A-M, Nikic S, Picault N et al (2000) *Arabidopsis SGS2* and *SGS3* genes are required for posttranscriptional gene silencing and natural virus resistance. Cell 101: 533–542

80. Nakanishi K, Weinberg DE, Bartel DP, Patel DJ (2012) Structure of yeast Argonaute with guide RNA. Nature 486: 368–374

81. Pall GS, Codony-Servat C, Byrne J, Ritchie L, Hamilton A (2007) Carbodiimide-mediated cross-linking of RNA to nylon membranes improves the detection of siRNA, miRNA and piRNA by northern blot. Nucleic acids research 35: e60

82. Parker JS, Roe SM, Barford D (2005) Structural insights into mRNA recognition from a PIWI domain–siRNA guide complex. Nature 434: 663–666

83. Peragine A, Yoshikawa M, Wu G, Albrecht HL, Poethig RS (2004) SGS3 and SGS2/SDE1/RDR6 are required for juvenile development and the production of trans-acting siRNAs in Arabidopsis. Genes & development 18: 2368–2379

84. Petrosius V, Aragon-Fernandez P, Üresin N, Kovacs G, Phlairaharn T, Furtwängler B, Op De Beeck J, Skovbakke SL, Goletz S, Thomsen SF et al (2023) Exploration of cell state heterogeneity using single-cell proteomics through sensitivity-tailored data-independent acquisition. Nature communications 14: 5910

85. Pfaff J, Hennig J, Herzog F, Aebersold R, Sattler M, Niessing D, Meister G (2013) Structural features of Argonaute-GW182 protein interactions. Proceedings of the National Academy of Sciences of the United States of America 110: E3770–3779

86. Poulsen C, Vaucheret H, Brodersen P (2013) Lessons on RNA silencing mechanisms in plants from eukaryotic argonaute structures. The Plant cell 25: 22–37

87. Qi Y, Denli AM, Hannon GJ (2005) Biochemical specialization within Arabidopsis RNA silencing pathways. Molecular cell 19: 421–428

88. Rivas FV, Tolia NH, Song JJ, Aragon JP, Liu J, Hannon GJ, Joshua-Tor L (2005) Purified Argonaute2 and an siRNA form recombinant human RISC. Nature structural & molecular biology 12: 340–349

89. Robinson JT, Thorvaldsdóttir H, Winckler W, Guttman M, Lander ES, Getz G, Mesirov JP (2011) Integrative genomics viewer. Nature biotechnology 29: 24–26

90. Sakurai Y, Baeg K, Lam AYW, Shoji K, Tomari Y, Iwakawa H-o (2021) Cell-free reconstitution reveals the molecular mechanisms for the initiation of secondary siRNA biogenesis in plants. Proceedings of the National Academy of Sciences 118: e2102889118

91. Schirle NT, MacRae IJ (2012) The crystal structure of human Argonaute2. Science 336: 1037–1040

92. Sheu-Gruttadauria J, MacRae IJ (2018) Phase Transitions in the Assembly and Function of Human miRISC. Cell 173: 946–957.e916

93. Shukla A, Yan J, Pagano DJ, Dodson AE, Fei Y, Gorham J, Seidman JG, Wickens M, Kennedy S (2020) poly(UG)-tailed RNAs in genome protection and epigenetic inheritance. Nature 582: 283–288

94. Sievers F, Wilm A, Dineen D, Gibson TJ, Karplus K, Li W, Lopez R, McWilliam H, Remmert M, Söding J et al (2011) Fast, scalable generation of high-quality protein multiple sequence alignments using Clustal Omega. Molecular Systems Biology 7: 539

95. Song JJ, Smith SK, Hannon GJ, Joshua-Tor L (2004) Crystal structure of Argonaute and its implications for RISC slicer activity. Science 305: 1434–1437

96. Souret FF, Kastenmayer JP, Green PJ (2004) AtXRN4 degrades mRNA in Arabidopsis and its substrates include selected miRNA targets. Molecular cell 15: 173–183

97. Takeda A, Iwasaki S, Watanabe T, Utsumi M, Watanabe Y (2008) The mechanism selecting the guide strand from small RNA duplexes is different among argonaute proteins. Plant & cell physiology 49: 493–500

98. Tan H, Luo W, Yan W, Liu J, Aizezi Y, Cui R, Tian R, Ma J, Guo H (2023) Phase separation of SGS3 drives siRNA body formation and promotes endogenous gene silencing. Cell reports 42

99. Till S, Lejeune E, Thermann R, Bortfeld M, Hothorn M, Enderle D, Heinrich C, Hentze MW, Ladurner AG (2007) A conserved motif in Argonaute-interacting proteins mediates functional interactions through the Argonaute PIWI domain. Nature structural & molecular biology 14: 897–903

100. Tsutsui H, Higashiyama T (2017) pKAMA-ITACHI Vectors for Highly Efficient CRISPR/Cas9-Mediated Gene Knockout in Arabidopsis thaliana. Plant and Cell Physiology 58: 46–56

101. Vaucheret H (2008) Plant ARGONAUTES. Trends in plant science 13: 350–358

102. Vazquez F, Vaucheret H, Rajagopalan R, Lepers C, Gasciolli V, Mallory AC, Hilbert JL, Bartel DP, Crete P (2004) Endogenous trans-acting siRNAs regulate the accumulation of Arabidopsis mRNAs. Molecular cell 16: 69–79

103. Vella MC, Choi EY, Lin SY, Reinert K, Slack FJ (2004) The C. elegans microRNA let-7 binds to imperfect let-7 complementary sites from the lin-41 3’UTR. Genes & development 18: 132–137

104. Vigh ML, Bressendorff S, Thieffry A, Arribas-Hernández L, Brodersen P (2022) Nuclear and cytoplasmic RNA exosomes and PELOTA1 prevent miRNA-induced secondary siRNA production in Arabidopsis. Nucleic acids research 50: 1396–1415

105. Vigh ML, Thieffry A, Arribas-Hernández L, Brodersen P (2024) Arabidopsis terminal nucleotidyl transferases govern secondary siRNA production at distinct steps. bioRxiv: 2024.2005.2027.596008

106. Wallmann A, Van de Pette M (2024) Structural and evolutionary determinants of Argonaute function. bioRxiv: 2024.2010.2010.617571

107. Wang T, Zheng Y, Tang Q, Zhong S, Su W, Zheng B (2021) Brassinosteroids inhibit miRNA-mediated translational repression by decreasing AGO1 on the endoplasmic reticulum. Journal of Integrative Plant Biology 63: 1475–1490

108. Wang XB, Jovel J, Udomporn P, Wang Y, Wu Q, Li WX, Gasciolli V, Vaucheret H, Ding SW (2011) The 21-nucleotide, but not 22-nucleotide, viral secondary small interfering RNAs direct potent antiviral defense by two cooperative argonautes in Arabidopsis thaliana. The Plant cell 23: 1625–1638

109. Wang Y, Juranek S, Li H, Sheng G, Wardle GS, Tuschl T, Patel DJ (2009) Nucleation, propagation and cleavage of target RNAs in Ago silencing complexes. Nature 461: 754–761

110. Wu P-H, Isaji M, Carthew Richard W (2013) Functionally Diverse MicroRNA Effector Complexes Are Regulated by Extracellular Signaling. Molecular cell 52: 113–123

111. Xiao Y, Maeda S, Otomo T, MacRae IJ (2023) Structural basis for RNA slicing by a plant Argonaute. Nature structural & molecular biology 30: 778–784

112. Xie Z, Allen E, Wilken A, Carrington JC (2005) DICER-LIKE 4 functions in trans-acting small interfering RNA biogenesis and vegetative phase change in Arabidopsis thaliana. Proceedings of the National Academy of Sciences of the United States of America 102: 12984–12989

113. Yang L, Wu G, Poethig RS (2012) Mutations in the GW-repeat protein SUO reveal a developmental function for microRNA-mediated translational repression in Arabidopsis. Proceedings of the National Academy of Sciences of the United States of America 109: 315–320

114. Yoshikawa M, Han Y-W, Fujii H, Aizawa S, Nishino T, Ishikawa M (2021) Cooperative recruitment of RDR6 by SGS3 and SDE5 during small interfering RNA amplification in Arabidopsis. Proceedings of the National Academy of Sciences 118: e2102885118

115. Yoshikawa M, Iki T, Numa H, Miyashita K, Meshi T, Ishikawa M (2016) A Short Open Reading Frame Encompassing the MicroRNA173 Target Site Plays a Role in trans-Acting Small Interfering RNA Biogenesis. Plant physiology 171: 359–368

116. Yoshikawa M, Iki T, Tsutsui Y, Miyashita K, Poethig RS, Habu Y, Ishikawa M (2013) 3’ fragment of miR173-programmed RISC-cleaved RNA is protected from degradation in a complex with RISC and SGS3. Proceedings of the National Academy of Sciences of the United States of America 110: 4117–4122

117. Yoshikawa M, Peragine A, Park MY, Poethig RS (2005) A pathway for the biogenesis of trans-acting siRNAs in Arabidopsis. Genes & development 19: 2164–2175

118. Zamore PD, Haley B (2005) Ribo-gnome: the big world of small RNAs. Science 309: 1519–1524

119. Zhang C, Ng DW-K, Lu J, Chen ZJ (2012) Roles of target site location and sequence complementarity in trans-acting siRNA formation in Arabidopsis. The Plant Journal 69: 217–226

120. Zhang X, Smits AH, van Tilburg GBA, Ovaa H, Huber W, Vermeulen M (2018) Proteome-wide identification of ubiquitin interactions using UbIA-MS. Nature protocols 13: 530–550

