## Supplementary figures and images for "The tryptophan-binding pockets of Arabidopsis AGO1 facilitate amplified RNA interference via SGS3"

### Appendix Figure 1

#### Figure 3A

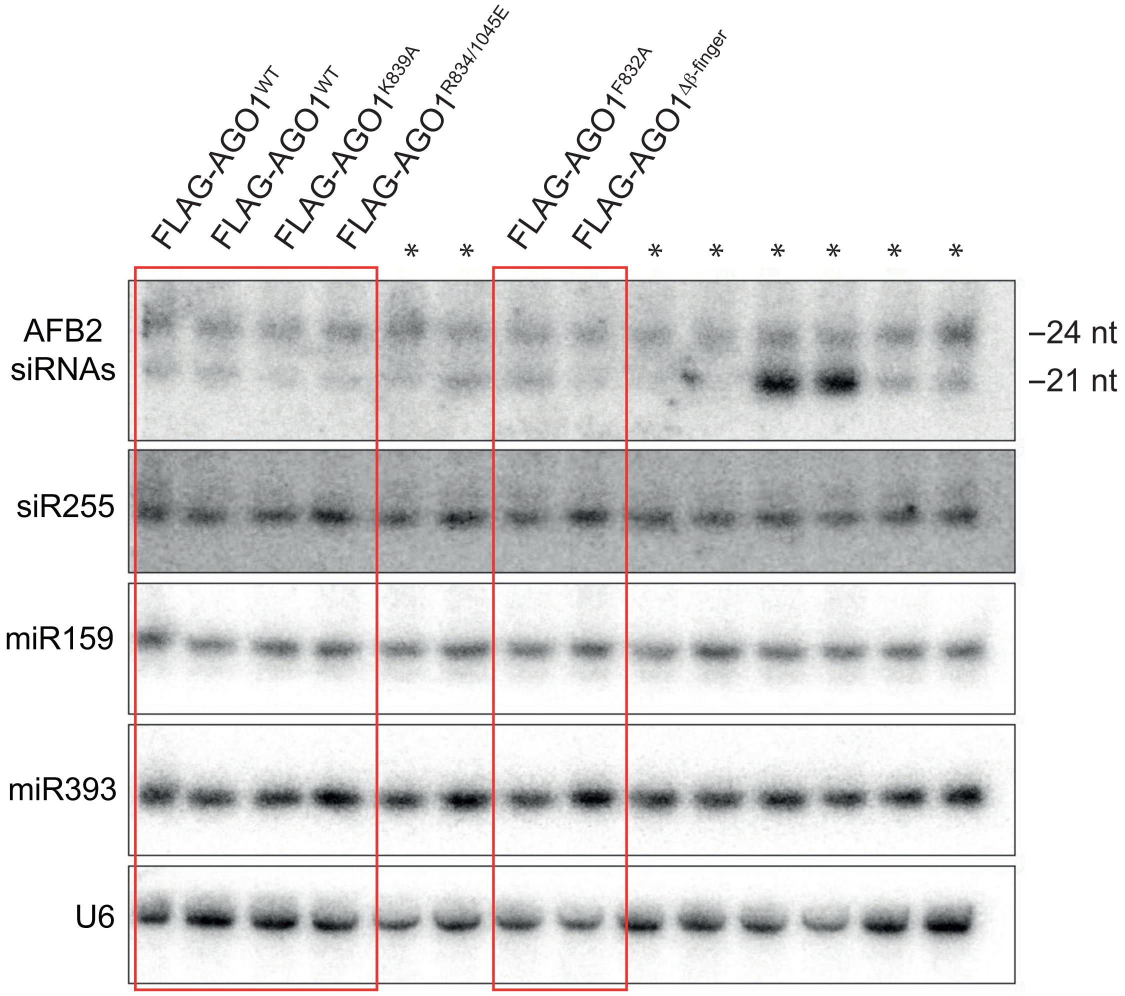


*Not relevant for this study

### Appendix Figure 2

#### Figure 6B

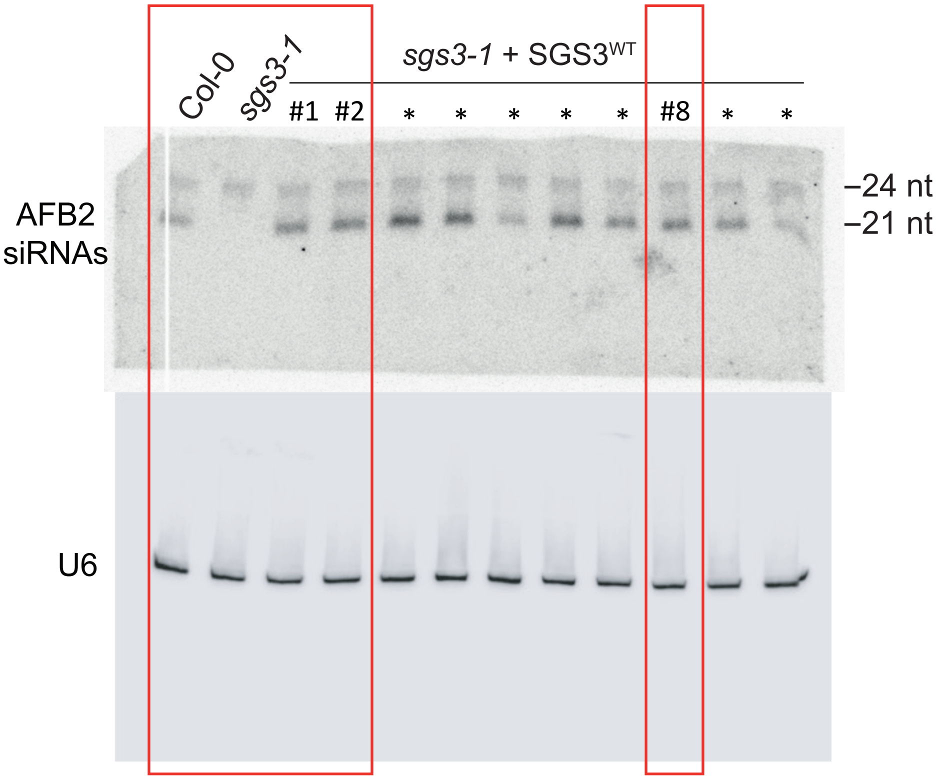


*Other transgenic lines
